# Long-lasting tagging of neurons activated by seizures or cocaine administration in Egr1-CreER^T2^ transgenic mice

**DOI:** 10.1101/2020.10.12.335745

**Authors:** Sophie Longueville, Yuki Nakamura, Karen Brami-Cherrier, Renata Coura, Denis Hervé, Jean-Antoine Girault

## Abstract

Permanent tagging of neuronal ensembles activated in specific experimental situations is an important objective to study their properties and adaptations. In the context of learning and memory these neurons are referred to as engram neurons. Here we describe and characterize a novel mouse line, Egr1-CreER^T2^, which carries a transgene in which the promoter of the immediate early gene *Egr1* drives the expression of the CreER^T2^ recombinase that is only active in the presence of tamoxifen metabolite, 4-hydroxy-tamoxifen (4-OHT). Egr1-CreER^T2^ mice were crossed with various reporter mice, Cre-dependently expressing a fluorescent protein. Without tamoxifen or 4-OHT, no or few tagged neurons were observed. Epileptic seizures induced by pilocarpine or pentylenetetrazol in the presence of tamoxifen or 4-OHT, induced the persistent tagging of many neurons and some astrocytes in the dentate gyrus of hippocampus, where *Egr1* is transiently induced by seizures. One week after cocaine and 4-OHT administration, these mice displayed a higher number of tagged neurons in the dorsal striatum than saline/4-OHT controls, with differences between reporter lines. Cocaine-induced tagging required ERK activation and tagged neurons were more likely than others to exhibit ERK phosphorylation or Fos induction after a second injection. Interestingly neurons tagged in saline-treated mice also had an increased propensity to express Fos, suggesting the existence of highly responsive striatal neurons susceptible to be re-activated by cocaine repeated administration, which may contribute to the behavioral adaptations. Our report validates a novel transgenic mouse model for permanently tagging activated neurons and studying long term alterations of *Egr1*-expressing cells.

## Introduction

In response to environmental stimuli, the striatum integrates sensory-motor information from the cortex and thalamus with a complex reward-related signal, provided by the dopamine inputs (Schultz *et al*., 2017). This integrative process is thought to create a spatio-temporal pattern of discharge activity in a specific ensemble of striatal neurons. A common view assumes that this pattern of activity durably modifies the synaptic connections allowing procedural memory storage and learning through the creation of preferential neural networks. These processes promote the subsequent selection of behaviors with positive outcome or, in contrary, the avoidance of those with negative consequence (Mink, 1996; Redgrave *et al*., 1999). In order to study the mechanisms implicated, it is essential to identify and long-lastingly follow neurons involved in the transiently activated network. It is now well established that gene induction and protein synthesis in the activated neurons are essential steps in the long-term maintenance of brain plasticity and memory (Dragunow, 1996) and in these processes, the induction of immediate early genes (IEGs) within minutes following neuron activation plays a critical role. These genes produce proteins that directly affect synaptic functions or, alternatively, transcription factors that act as master genes regulating synapses. To study the ensembles of neurons selectively modified by specific events and presumably contributing to their memory traces, several strategies have been used, aiming to visualize elusive memory engrams, as proposed by Richard Semon at the beginning of the XX^th^ century. Several strategies have been developed over the last years, all based on the use of IEGs. The group of Tonegawa used a combination of *Fos* IEG promoter driving the tetracyclin-dependent activator (Fos-tTA) targeting tetracyclin-response element-dependent reporter genes (Liu *et al*., 2012) generating insightful results on hippocampus- and amygdala-dependent memory (review in (Tonegawa *et al*., 2015)). Others have used IEG promoters to induce a pharmacologically controlled form of the Cre recombinase (CreER^T2^) inducing recombination of LoxP-dependent reporter genes (Guenthner *et al*., 2013; DeNardo & Luo, 2017). The permanent recombination events in activated neurons occur during a time window defined by tamoxifen administration. Until today, the promoters of Fos and Arc/Arg3.1 have been used in different lines of transgenic mice and recombinant viruses (Guenthner *et al*., 2013; Denny *et al*., 2014; Ye *et al*., 2016; DeNardo *et al*., 2019). The development of these genetic tools has led to very interesting studies shedding new insights in a wide variety of neural processes (Ye *et al*., 2016; Allen *et al*., 2017; Girasole *et al*., 2018; Sanders *et al*., 2019).

We independently developed a mouse model using CreER^T2^ driven by the promoter of *Egr1* (also known as *Zif268*, *Krox24*, *TIS-8* and *NGF1-A*). We chose this gene because it is crucial for certain forms of long-term memory (Walton *et al*., 1999; Bozon *et al*., 2003a; Bozon *et al*., 2003b). *Egr1* is expressed in basal conditions in several brain regions, including the striatum (Herdegen *et al*., 1995), or in response to natural or pharmacological stimuli in specific neural networks (Worley *et al*., 1991; Beckmann & Wilce, 1997). The basal expression of *Egr1* is tightly linked to physiological firing activity of neurons, since it is strongly reduced by AMPA or NMDA glutamatergic receptor antagonists as well as by L-type voltage-sensitive calcium channel blockers (Murphy *et al*., 1991; Vaccarino *et al*., 1992; Wang *et al*., 1994). Psychostimulants or reward learning cause induction of *Egr1* in striatal neurons (Cole *et al*., 1992; Moratalla *et al*., 1992; Bhat & Baraban, 1993; Maroteaux *et al*., 2014). Invalidation of the *Egr1* gene has no effect on the acute effects of cocaine, but greatly reduces the sensitization to repeated cocaine administrations and abolishes their rewarding effects (Valjent *et al*., 2006). Likewise, this invalidation reduces the food operant conditioning (Maroteaux *et al*., 2014). These studies suggest that *Egr1* is involved in neuronal plasticity that remodels activated neural networks to consolidate stable memory traces (Veyrac *et al*., 2014). The cells in which *Egr1* is induced are likely to be those that undergo stable synaptic changes, leading to durable responses to cocaine or sustained instrumental behaviors to earn food.

In the present study we describe a novel mouse line, designated as Egr1-CreER^T2^, carrying a bacterial artificial chromosome (BAC) transgene in which the *Egr1* promoter controls the expression of CreER^T2^. First, we characterized this line and show that it allows permanent labeling of activated neurons, using epilepsy models known to strongly induce *Egr1* in the hippocampus (Sukhatme *et al*., 1988; Hughes & Dragunow, 1994; Beckmann & Wilce, 1997; Li *et al*., 2005). We then explored the effects of intense stimulation of reward circuit following cocaine administration. By crossing the Egr1-CreER^T2^ mice with several reporter mice, we found a higher number of tagged neurons in the dorsal striatum after cocaine administration than after saline. However, the basal and cocaine-evoked tagging varied depending on the reporter mice used. Cocaine-induced tagging required ERK activation as predicted by the role of ERK activation in the induction of *Egr1* by cocaine (Valjent *et al*., 2006). We also observed that the population of labeled neurons was more activated by a new cocaine injection than unlabeled neurons. Our study shows that this novel Egr1-CreER^T2^ system is a valuable tool to permanently tag neurons whose activation includes *Egr1* induction. In the striatum it identifies neurons activated by cocaine, and potentially other rewarding stimuli. It will be helpful for studying long-term physiological or morphological evolution of the cocaine-activated network and its role in striatal-linked behaviors or other types of long-term neuronal modifications in diverse brain regions.

## Materials and Methods

### Animals

The Tg(Egr1-CreER^T2^) mouse line (here referred to as Egr1-CreER^T2^) was generated in 2009 at the Mouse Clinical Institute (MCI, Strasbourg, France). A bacterial artificial chromosome (BAC, clone RP23-70M6) that contained the entire mouse *Egr1* gene with 95 kb 5’ upstream and 86 kb downstream regions was used. This BAC was modified by homologous recombination by transfecting bacteria with a DNA fragment containing a cassette encoding a fusion protein (CreER^T2^) consisting of a Cre recombinase (Cre) and a mutated ligand-binding domain of the human estrogen receptor (ERT2) (Feil *et al*., 1997) as well as two homology arms of *Egr1* gene regions 3’ and 5’ from the transcription start and a neomycin resistance gene flanked by FRT sequences. The recombined BAC was transfected into mouse embryonic stem (ES) cells to obtain neomycin-resistant transgenic cells. The neomycine resistance gene was then removed by flipase and the resulting transgenic ES cells was injected into C57Bl/6J blastocysts to obtain chimeric mice. Germ line transmission of the transgene was obtained for two lines. Since they provided similar results in the first pilot explorations, the subsequent experiments were performed using only one mouse line.

The Egr1-CreER^T2^ mice were kept heterozygous on a C57Bl/6J genetic background. They were crossed with four reporter mouse lines with recombined Rosa26 locus, in which various fluorescent proteins can be expressed following Cre-dependent recombination: R26^RCE^ (Gt(ROSA)26Sor^tm1.1(CAG-EGFP)Fsh^) (gift of Dr G Fishell) (Miyoshi *et al*., 2010), R26^Ai14^ (Gt(ROSA)26Sor^tm14(CAG-tdTomato)Hze^) (gift of Dr JC Poncer) (Madisen *et al*., 2010), R26^Trap^ (Gt(ROSA)26Sor^tm9(EGFP/Rpl10a)Amc^) (Liu *et al*., 2014) and R26^hM4D^ (Gt(ROSA)26Sor^tm1(CAG-CHRM4*,-mCitrine)Ute^)/J (Zhu *et al*., 2016). The last two lines were purchased from the Jackson Laboratory (Bar Harbor, USA). The various Rosa26-recombined mice were kept homozygous and crossed with heterozygous Egr1-CreER^T2^ mice.

Egr1-CreER^T2 Tg/+^ mice were crossed by *Drd1*-EGFP ^Tg/+^ (Tg(Drd1-EGFP)X60Gsat/Mmmh, RRID:MMRRC_000297-MU) or Drd2-L10aEGFP ^Tg/+^ (Tg(Drd2-EGFP/Rpl10a)CP101Htz/J) mice (gift of Pr P Greengard) that express EGFP and EGFP-Rpl10a ribosomal fusion protein in the neurons that express D1 and D2 dopamine receptors (Drd1 and Drd2), respectively. The resulting doubly transgenic mice were used for identifying whether the activity-tagged neurons express Drd1 or Drd2 in the striatum.

The mice were maintained in a 12 h light/dark cycle in stable conditions of temperature (22°C) with access to food and water ad libitum. All animal procedures used in the present study were approved by the Ethical committee for animal experiments (*Comité d’Ethique en Expérimentation Animale Charles Darwin*, Paris, France) and by the *Ministère de l’Education Nationale de l’Enseignement Supérieur de la Recherche* (France, project APAFIS#07636.03). All methods in this study were performed in accordance with the relevant guidelines and regulations

### Drugs

Tamoxifen (T5648, Sigma-Aldrich) and 4-hydroxytamoxifen (H7904 or H6278, Sigma-Aldrich) were dissolved in 1 mL ethanol (100%). Corn oil (C8267, Sigma-Aldrich) was then added and ethanol was evaporated by incubating the open tubes at 50°C overnight in the dark. The volume of corn oil was adjusted for injecting the dose of active drug at 10 mL.kg^−1^. Tamoxifen (from 50 to 220 mg.kg^−1^) was administered i.p. and occasionally by gavage, using a feeding cannula adapted for mice. 4-hydroxytamoxifen (4-OHT) was administered by i.p. injection at the dose of 50 mg.kg^−1^ of active Z form. Lithium chloride (L4408, Sigma-Aldrich, 10.6 mL.kg^−1^), methylscopolamine (S2250, Sigma-Aldrich - 10 ml.kg^−1^), pilocarpine (P6503, Sigma-Aldrich, 7 mL.kg^−1^), cocaine HCl (Cooper France, 10 mL.kg^−1^), and pentylenetetrazol (P6500, Sigma-Aldrich, 7.5 mL.kg^−1^) were dissolved in saline solution (NaCl 0.9 g.L^−1^) and injected i.p. α-[amino-[(4-aminophenyl)thio]methylene]-2-(trifluoromethyl)-benzene-acetonitrile (SL327; Sigma-Aldrich) was dissolved in 5% DMSO, 5% Tween 80, 15% polyethyleneglycol 400 (all vol per vol) in water and injected i.p. (5 mL.kg^−1^). Diazepam TVM 5 (Laboratoire TVM, France) was injected i.p. at 4 mL.kg^−1^.

### Stereotaxic injection of viral vectors

Cre-inducible adeno-associated virus (AAV) vectors coding for mCherry, pAAV8-hSyn-DIO-mCherry (Addgene, Cambridge, MA), were bilaterally injected into the dorsal striatum of doubly transgenic mice bearing the Egr1-CreER^T2^ transgene in association with Drd1-EGFP or Drd2-Rpl10aEGFP transgene. Animals were anesthetized with a mix of ketamine (80 mg.kg^−1^) and xylazine (10 mg.kg^−1^) and mounted on a stereotaxic apparatus (Kopf, France). Two injections (0.5 μl, each), were performed dorsal striatum, one in the right and one in the left side, at the following coordinates based on the Allen Brain Atlas: anteroposterior 0.1 mm, mediolateral ± 2.3 mm, dorsoventral −3.75 from the bregma. The injections were performed using a 10 μl Hamilton Neuros 33G syringe at a flow rate of 0.2 μl.min^−1^. The needle which express was left in place for 2 min before and 5 min after the injection and then slowly retracted. Mice were placed in their home cage for 2 weeks for recovery.

### Induction of epilepsy

Pilocarpine treatment was used for status epilepticus induction (Curia *et al*., 2008). Four-month old Egr1-CreER^T2 Tg/+^ x R26^RCE/+^ mice (n = 33) were pretreated with LiCl (423 mg.kg^−1^, i.p.) and methylscopolamine (1 mg.kg^−1^, i.p.), 24 h and 30 min before pilocarpine, respectively, to increase the central effects of pilocarpine and reduce its peripheral effects. Pilocarpine (70 mg.kg^−1^, i.p.) was administered at the same time as tamoxifen (120 mg.kg^−1^, i.p.). If no seizure was observed after 1 h, the mice were injected again with pilocarpine (50 mg.kg^−1^, i.p.). In 76% of the mice (n = 25), intermittent seizures appeared 40-120 min after the pilocarpine injection and were followed at 50-130 min by a *status epilepticus* characterized by generalized tonic-clonic seizures. This *status epilepticus* was blocked 45-60 min later by an administration of diazepam (10 mg.kg^−1^). About 24% of the mice (n = 8) did not develop any apparent seizures. Control mice (n = 3) received LiCl, methylscopolamine and tamoxifen but not pilocarpine. During the next 3 days, the mice were kept in warm atmosphere by placing their cages on a warm plate (30°C). They received daily injections of 5% glucose in saline (0.3 ml, s.c.) and were fed with enriched food. Despite this care, about 19% of the epileptic mice died following pilocarpine treatment. Animals were sacrificed by paraformaldehyde perfusion (see below) at various times after these treatments (ranging from 24 h to 1 month). In one experiment, tamoxifen was injected 7 weeks after the pilocarpine treatment, when pilocarpine-treated mice (n = 6) displayed intermittent seizures.

We also induced epileptic seizures in three-month old Egr1-CreER^T2 Tg/+^ x R26^Ai14/+^ (n = 3) or CreER^T2 Tg/+^ x R26^Trap/+^ (n = 2) mice by treating them with pentylenetetrazol (PTZ, 37.5 mg.kg^−1^, s.c.). This treatment was associated with an injection of the short-acting tamoxifen metabolite, 4-hydroxytamoxifen (4-OHT, 50 mg.kg^−1^, i.p.). In the minutes following the PTZ injection, general convulsions usually appeared. The animals recovered a normal behavior after 2 h.

### Cocaine, 4-hydroxytamoxifen treatment and locomotor activity quantification

Locomotor activity was recorded by automated video-tracking in a 0.5 m wide square arena (VideoTrack, ViewPoint, Lyons, France). During the 12 days preceding the cocaine and 4-OHT treatments, Egr1-CreER^T2 Tg/+^ x R26^RCE/+^, Egr1-CreER^T2 Tg/+^ x R26^Ai14/+^ or Egr1-CreER^T2 Tg/+^ x R26^Trap/+^ mice (age range between 2 and 6 months) were habituated to the experimental conditions. They were handled the first 3 days, 5 min per day. Then, the next 2 days, they were handled and pricked in the abdomen with a syringe needle without injection. The next 4 days, mice were handled and administered with saline (3 days, 10 mL.kg-1, and 1 day, 20 mL.kg-1, i.p.). Then, during last 2 days of habituation, mice were placed 2 h in the arena, and injected with saline (20 mL.kg^−1^) after 1h. On the day of the experiment, the animals were placed in the VideoTrack arena and 1 h later received a 4-OHT injection (50 mg.kg^−1^, i.p.), immediately followed by cocaine (20 mg.kg^−1^, i.p. in 0.9 g.L^−1^ NaCl) or saline injection. After the injections, mice were returned to the arena where they stayed 3 additional hours. The distance travelled was analyzed on the video using the VideoTrack software. Most animals were perfused with fixative one week later (see below). In some Egr1-CreER^T2 Tg/+^ x R26^Ai14/+^ mice, a second injection of cocaine was performed 7 days after the cocaine or saline treatment and the locomotor activity was recorded under conditions identical to those for the first injection, except the animals were perfused with fixative solution 15 min or 90 min after cocaine injection.

### Immunofluorescence

Mice were rapidly and deeply anesthetized with pentobarbital (500 mg.kg^−1^, i.p., Dolethal, Vetoquinol, France) and perfused transcardially with 40 g.L^−1^ paraformaldehyde in 0.1 M sodium phosphate buffer pH 7.4. Brains were post-fixed overnight at 4°C and cut into 30 μm-thick free-floating sections with a vibrating microtome (Leica). Brain sections were kept at −20°C in a solution containing 300 mL.L^−1^ ethylene glycol, 300 mL.L^−1^ glycerol and 0.1 M phosphate buffer. Immunolabeling procedures were as previously described (Valjent *et al*., 2000). After three 10-min washes in Tris-buffered saline (TBS, 0.10 M Tris and 0.14 M NaCl, pH 7.4), free floating brain sections were incubated for 5 min in TBS containing 30 mL.L^−1^ H_2_O_2_ and 100 mL.L^−1^ methanol, and rinsed again in TBS 3 times for 10 min. Brain sections were then incubated for 15 min in 2 mL.L^−1^ Triton X-100 in TBS, rinsed 3 times in TBS and were blocked in 30 g.L^−1^ bovine serum albumin in TBS, and then incubated overnight at 4°C with the primary antibody diluted in the same blocking solution. Sections were then washed three times in TBS for 15 min and incubated 1 h with secondary antibodies. After washing again, sections were mounted in Vectashield (Vector laboratories).

Primary antibodies were mouse antibodies against calbindin (#300, Swant), glial fibrillary acidic protein (GFAP, 1:1000, #SAB4100002, Sigma-Aldrich), NeuN (1:500, #MAB377, Millipore), parvalbumin (1:1000, #PV235, Swant), rabbit antibodies against KCC2 (1:1000, #7432, Millipore), Cre recombinase (a gift of F. Tronche, 1:1000,), Egr1 (1:2000, #4154, Cell Signaling), phospho-ERK1/2 (Thr202/204) (1:1000, #9101, Cell Signaling), Fos (1:1000, #sc-52, Santa Cruz), and chicken antibodies against mCherry (1:1500; Abcam, #ab205402 RRID:AB_2722769). Secondary antibodies were anti-rabbit A488 antibody (1:400, Invitrogen, #A21206), anti-rabbit Cy3 antibody (1:400; Jackson Immuno Research, #711-165-152), anti-mouse A488 antibody (1:400, Invitrogen, #A21202), anti-mouse Cy3 antibody (1:400; Jackson Immuno Research, #715-165-150), and anti-chicken Cy3 antibody (1:800; Jackson Immuno Research, #703-165-155). Images were acquired using confocal microscope (Leica SP5) with a 40X objective.

### Neuron counting

Neuron counts were performed throughout the right and left dorsal striatum in one section per animal using ImageJ. Sections were selected at the same level in different animals of the same experiment, between 1.0 and 0.3 mm anterior to Bregma. In one experiment, neuron counting was carried out in the granular layer of one dentate gyrus of hippocampus. The level of sections counted was 1.8 mm posterior to Bregma. DAPI-positive nuclei of striatal projection neurons (SPNs, a.k.a. medium-size spiny neurons) were counted using a home-made ImageJ macro. Briefly, DAPI pictures were background corrected using the rolling ball method. A threshold was then chosen using the ‘Auto threshold’ function. Cells were subsequently evaluated in the whole dorsal striatum by running the ‘Analyze particles’ function in ImageJ. Based on size and shape criteria (see (Matamales *et al*., 2009)), SPN nuclei were identified by a mask of positive particles and their number evaluated. The activity-tagged neurons that were positive for EGFP, tdTomato or mCherry were quantified manually by an observer unaware of the mouse groups. The % of activity-tagged SPNs in the total number of DAPI-positive SPNs was evaluated in the left and right dorsal striatum or granular layer of dentate gyrus. To evaluate the number of pERK or Fos positive neurons, we applied DAPI-positive SPN masks on pERK or Fos images and we automatically quantified the % of pERK or Fos positive neurons in DAPI-positive SPN by defining a threshold of pERK or Fos labeling. In the populations of pERK- or Fos-positive neurons, we determined the number of activity-tagged and non-tagged neurons manually.

### Statistical analyses

The data were tested from deviation from normal distribution with D’Agostino & Pearson omnibus and Shapiro-Wilk normality tests. When deviation from normal distribution was not significant and number of samples ≥ 7 differences between two groups were analyzed with unpaired Student’s t test and between four groups with two-way ANOVA followed by Holmes-Sidak’s post hoc tests. Time course were analyzed with repeated measures ANOVA. When distribution was not normal or sample size small non parametric tests were used, Mann-Whitney’s test for two groups and Kruskal-Wallis for more groups followed by post-hoc Dunn’s test. All statistical analyses were performed using GraphPad Prism 6.07. A threshold value for significance of p < 0.05 was used throughout the study. The detailed results of statistical analyses are provided in **Supplementary Table 1**.

## Results

### Generation and characterization of Egr1-CreER^T2^ mice

To permanently tag neurons activated by a specific experience, we used the *Egr1* promoter to drive the expression of tamoxifen-inducible Cre recombinase (CreER^T2^) and to produce a recombination event in a Cre-dependent transgene or virus. With this approach, when a neuron is activated, the activation of Egr1 promoter induces CreER^T2^ production. CreER^T2^ is trapped in the cytoplasm, but if tamoxifen (or 4-hydroxytamoxifen, 4-OHT) is present, CreER^T2^ enters into the nucleus and recombines the transgene, resulting in its permanent expression (**Fig. 1A, B**). The new mouse line that we generated, Tg(Egr1^CreERT2^), referred to below as Egr1-CreER^T2^, carried a recombined BAC in which the coding sequence of the IEG gene, *Egr1*, was replaced by that of CreER^T2^. We first verified the expression of CreER^T2^ in transgenic mice and its nuclear accumulation in response to tamoxifen. We treated Egr1-CreER^T2^ mice with cocaine (20 mg.kg^−1^, i.p., **Fig. 1C**). Without tamoxifen, 5 h after cocaine we observed a low diffuse Cre immuno-labeling in both the cytoplasm and nucleus of striatal neurons (**Fig. 1C, upper panels**). In mice in which tamoxifen (165 mg.kg^−1^, p.o.) was administered 1 h before cocaine, we observed an accumulation of CreER^T2^ within the nucleus of striatal neurons (**Fig. 1C, lower panels**), indicating that tamoxifen efficiently stimulated the transport of CreERT^2^ into the nucleus.

**Figure 1:**
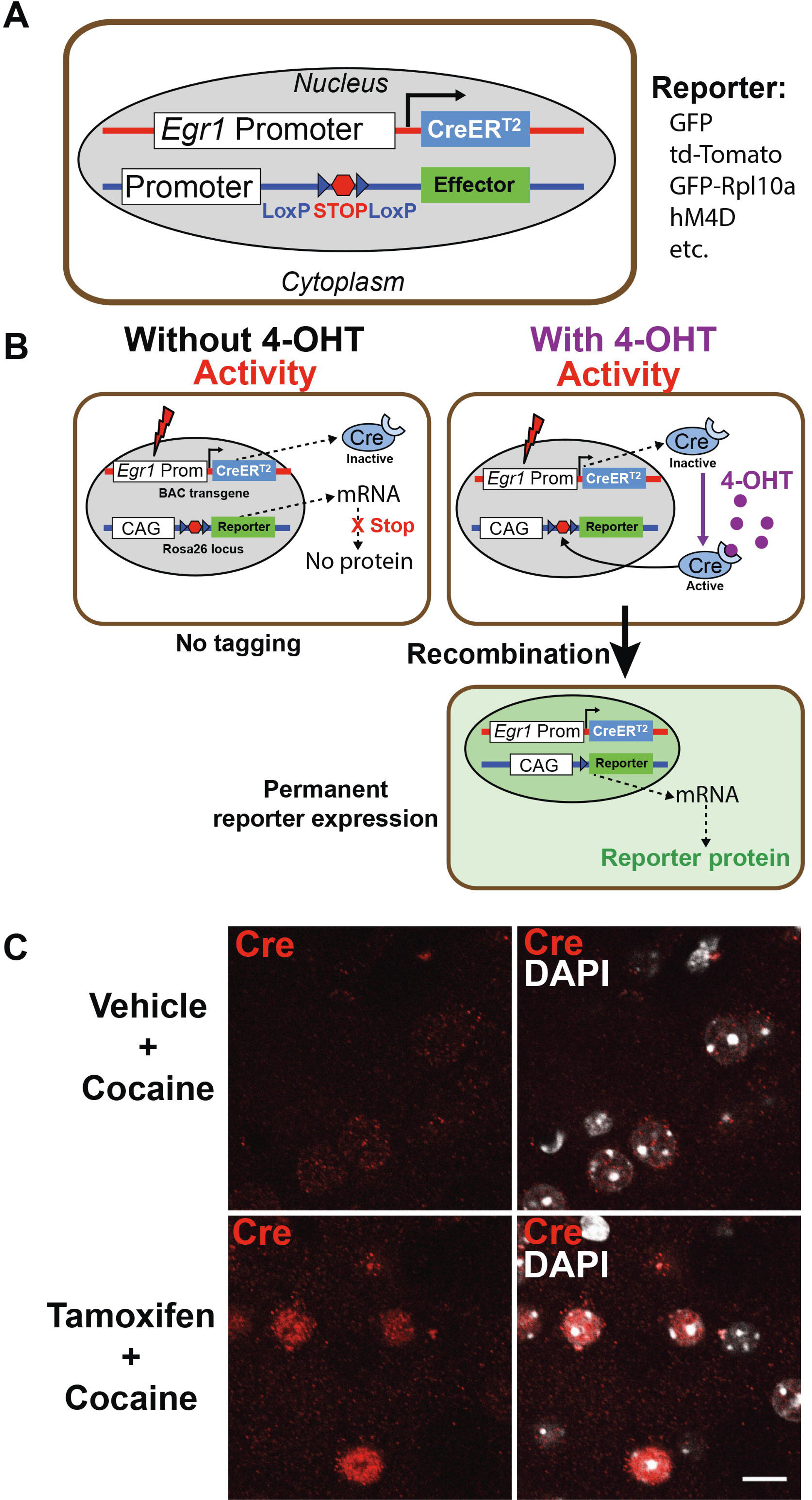
Strategy for permanent tagging of activated cells using the activity-dependent promoter of *Egr1*. **A**) Mice harbor two transgenes: one is a recombined BAC transgene expressing CreER^T2^ under the control of *Egr1* gene promoter and the other allows Cre-dependent expression of a fluorescent effector, such as EGFP (R26^RCE^), tdTomato (R26^Ai14^), EGFP fused with L10a ribosomal protein (Rpl10a) (R26^TRAP^) or hM4D fused with citrine (R26^hM4D^). **B**) When a neuron is stimulated, Egr1 promoter activation promotes the expression of CreER^T2^ protein. Without tamoxifen or its active metabolite, 4-hydroxytamoxifen (4-OHT), CreER^T2^ remains inactive in the cytoplasm preventing DNA recombination in the nucleus (left). The reporter gene including a strong promoter (e.g. derived from chicken actin globin, CAG) is transcribed but not translated because of the presence of a Stop codon. In contrast, the presence of 4-OHT elicits the nuclear translocation of active CreER^T2^, leading to the excision of the STOP codon and the expression of the effector protein. After this recombination event, the effector is permanently expressed. **C**) Egr1-CreER^T2 Tg/+^ mice were treated with cocaine (20 mg.kg^−1^) to stimulate *Egr1* promoter in striatal neurons. One hour later, mice were administered with tamoxifen (165 mg.kg^−1^, p.o.) or vehicle and perfused 4 h later. Brain sections were stained for Cre immunoreactivity (red) and DAPI (white). In vehicle-treated animals, CreER^T2^ immunoreactivity is poorly visible. In contrast, in tamoxifen-treated mice, CreER^T2^ accumulates in the nucleus of striatal neurons which become clearly immunoreactive. Scale bar: 10 µm.

We then examined whether in the absence of tamoxifen treatment there was a spontaneous recombination due to a leakage of Cre activity. We crossed the Egr1-CreER^T2^ mice with four lines with a recombined Rosa26 locus including EGFP (R26^RCE^) (Miyoshi *et al*., 2010), tdTomato (R26^Ai14^) (Madisen *et al*., 2010), Rpl10a-EGFP (R26^Trap^) (Liu *et al*., 2014) or hM4D receptor (R26^hM4D^) (Zhu *et al*., 2016). hM4D is a Gi-coupled designer receptor exclusively activated by designer drugs (DREADD). In the absence of tamoxifen, very few fluorescent neurons were observed within the striatum or hippocampus in the double transgenic mice (**Fig. 2A-D**). However, we noticed that their numbers differed between the various reporter lines used. The largest number of spontaneously positive cells was found in Egr1-CreER^T2 Tg/+^ x R26^Ai14/+^ mice, mainly in the cortex and striatum, less in the hippocampus (**Fig. 2A**). In these mice, tdTomato labelling could be also observed in axons and terminals in the thalamus. In contrast, the number of spontaneously recombined cells was extremely low in the striatum and hippocampus of the Egr1-CreER^T2 Tg/+^ x R26^RCE/+^ mice (**Fig. 2B**), and slightly higher in the Egr1-CreER^T2 Tg/+^ x R26^Trap/+^ and Egr1-CreER^T2 Tg/+^ x R26^hM4D/+^ mice (**Fig. 2C, D**). These data show that CreER^T2^ expression can produce recombination in few neurons in the absence of tamoxifen and that their number varies depending on the reporter mouse lines.

**Figure 2:**
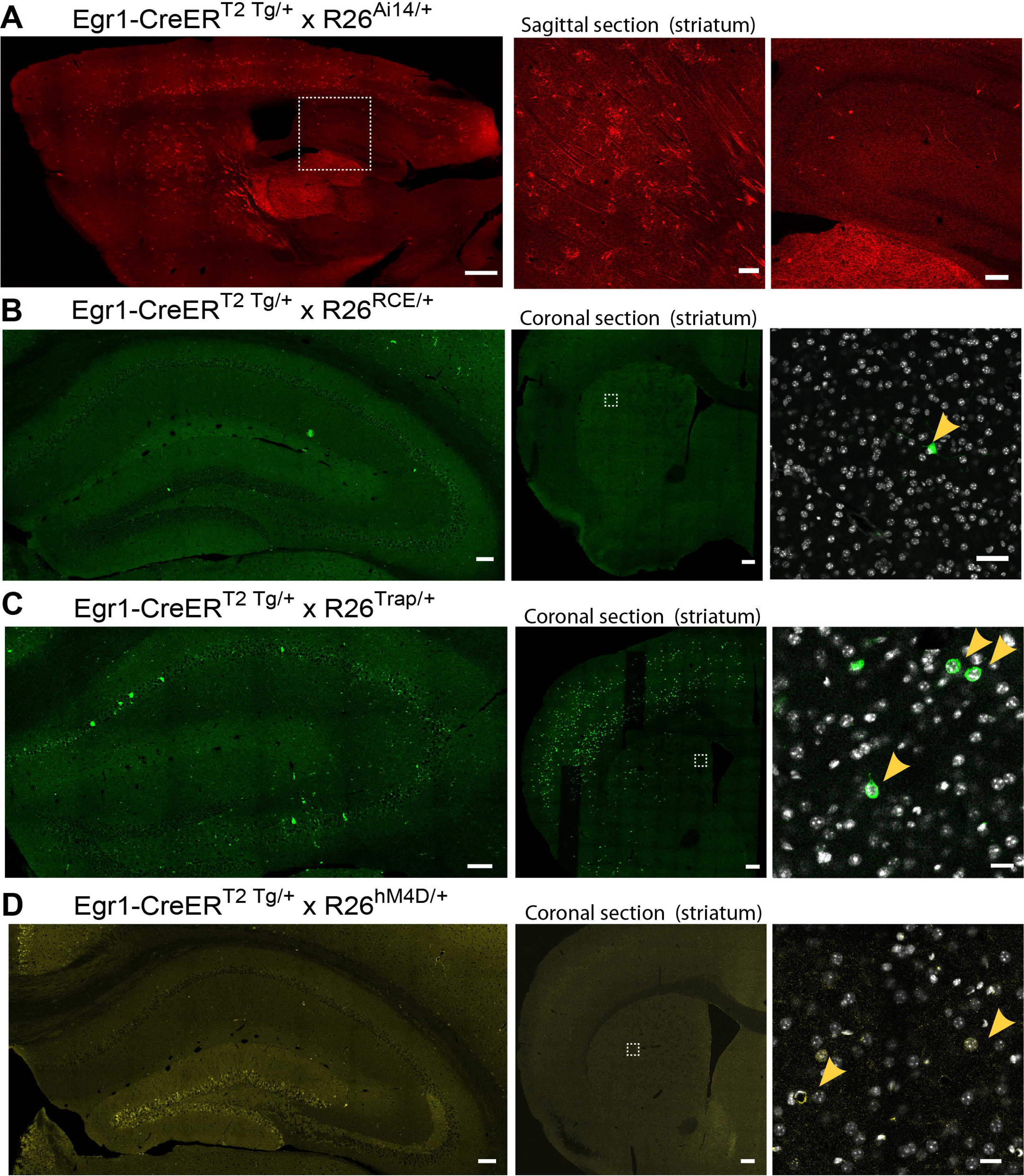
Recombination events are rare in the absence of tamoxifen treatment. Confocal sections showing the spontaneous fluorescence in Egr1-CreER^T2^ mice crossed with various reporter lines in which the transgene is inserted at the Rosa26 locus: R26^Ai14^ (**A**), R26^RCE^ (**B**), R26^Trap^ (**C**), and R26^h4MD^ (**D**) mice. These mice never received tamoxifen or 4-OHT. Left panels: sagittal whole brain section (**A**) or coronal sections through the hippocampus (**B, C, D**). Middle panels: hemisphere sections at the level of striatum. Right panels in **A**, higher magnification of the boxed area in left panel, in **B, C, D**, higher magnification of the boxed area in middle panel, showing rare cells with spontaneous recombination. The number of recombined cells varies depending on the reporter line: R26^Ai14/+^ >> R26^Trap/+^ = R26^hM4D/+^ > R26^RCE/+^ mice (all crossed with Egr1-CreER^T2 Tg/+^ mice). Scale bars: **A)** left 1 mm, middle and right 250 µm; **B)** left 100 µm, middle 250 µm, right 50 µm; **C-D)** left 100 µm, middle 250 µm, right 20 µm.

### Activity-dependent genetic labeling of hippocampal cells after pilocarpine-induced seizures

We then examined whether tamoxifen treatment was able to induce an efficient recombination in the target reporter gene after endogenous neuronal activation. It is well demonstrated that pilocarpine-induced epileptic seizures produce intense induction of Egr1 gene in the hippocampus, particularly in the dentate gyrus (DG) (Hughes & Dragunow, 1994; Lösing *et al*., 2017). To test the validity of the Egr1-CreER^T2^ system, we therefore investigated whether hippocampal neurons can be long-lastingly tagged following pilocarpine-induced *status epilepticus*. We treated Egr1-CreER^T2 Tg/+^ x R26^RCE/+^ mice tamoxifen (100 mg.kg^−1^) with or without pilocarpine and sacrificed them one week later to check the presence of EGFP-tagged neurons in the hippocampus. In control mice that had received tamoxifen but no pilocarpine, we observed no EGFP-positive cells in the DG or CA3, and only a few in CA1/CA2 area (**Fig. 3A, left panel**). In the pilocarpine- and tamoxifen-treated mice, the number of EGFP-tagged cells depended on whether the animals had displayed *status epilepticus* (**Fig. 3A, right panel**) or not (**Fig. 3A, middle panel**). The number of EGFP-positive cells was much higher in epileptic mice than in those without obvious epileptic seizure. Quantification of the number of EGFP-tagged cells in the granular layer of DG indicated that on average 6% of the neurons, with a maximum of 23%, were labeled in mice that had developed *status epilepticus* (**Fig. 3B**). In contrast, in mice without apparent epileptic seizure, their number averaged about 0.2%. These experiments showed that in tamoxifen-treated Egr1-CreERT2^Tg/+^ x R26^RCE/+^ mice neuronal activation during status epilepticus induced a recombination in a substantial number of neurons. However, in this mouse line and under these experimental conditions, only a relatively low proportion of the cells was tagged.

**Figure 3:**
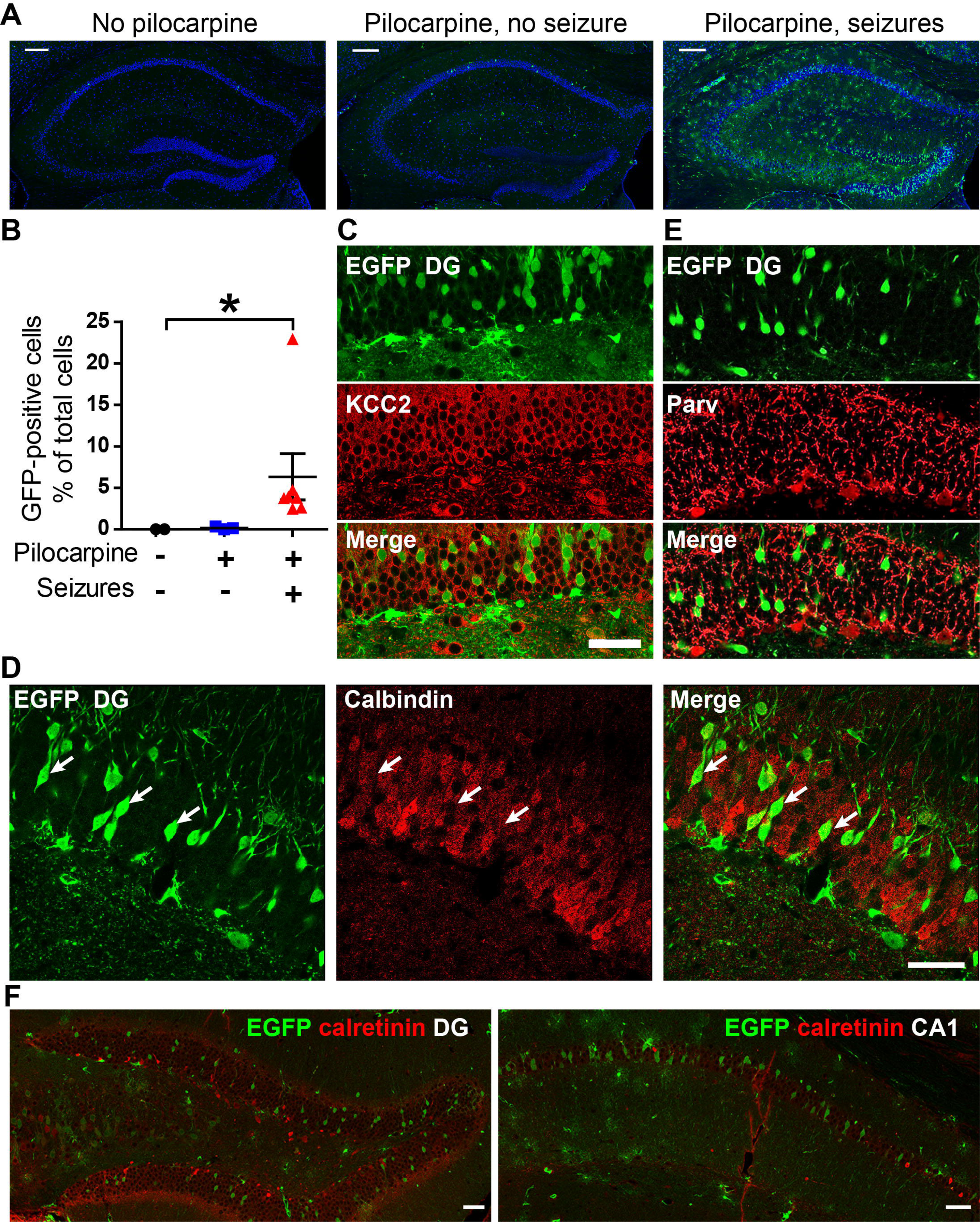
Persistent tagging of hippocampal neurons after pilocarpine-induced epileptic seizures in Egr1-CreER^T2^ mice. Egr1-CreER^T2 Tg/+^ x R26^RCE/+^ double transgenic mice (n = 9) were treated with pilocarpine (70 mg.kg^−1^, see methods) and immediately after tamoxifen (120 mg.kg^−1^). In two mice, pilocarpine was omitted and replaced by a saline injection. One week later, EGFP expression was investigated in the hippocampus. **A**) EGFP and DAPI fluorescence of coronal sections of hippocampus: *left*, mouse which did not receive pilocarpine, *middle,* pilocarpine-treated mouse that did not show any apparent seizure, *right*, pilocarpine-treated mouse that developed intense seizures. Scale bar: 200 µm **B**) Quantification of EGFP-positive neurons in the dentate gyrus (DG). Data are scatter plots with mean ± SEM (one point per mouse, n = 2, 3, and 7). Kruskal-Wallis, 8.45, p < 0.01 followed by Dunn’s multiple comparisons test, no pilocarpine vs pilocarpine seizures * p < 0.05 (See detailed statistical results in **Supplementary Table 1**). **C**) EGFP-positive neurons in the DG express high levels of KCC2 transporter immunofluorescence (red). **D**) EGFP activity-tagged neurons in the granular layer of hippocampus dentate gyrus express low levels of calbindin immunofluorescence (white arrows). Scale bar: 50µm. **E**) Interneurons immunoreactive for parvalbumin (Parv) were not EGFP-tagged in the DG. **F**) EGFP and calretinin immunofluorescence are not colocalized in the DG (left) and CA1 (right). Scale bar (**C-F**): 50 µm. Scale bar: 50 µm.

We then characterized the type of cells that were EGFP-tagged in epileptic animals. EGFP-tagged cells were found in the granular cell layer of the DG, the mossy cell region of hilus, and the pyramidal layer of CA3 and CA1. In the dentate gyrus, labelled neurons were granular cells that abundantly expressed the KCC2 neuronal protein (**Fig. 3C**) and low levels of calbindin (**Fig. 3D**). We did not observe any EGFP-labeling in GABAergic neurons expressing parvalbumin (**Fig. 3E**) or calretinin (**Fig. 3F**). No EGFP was found in microglial cells identified with Iba1 staining (**Fig. 4A**). In contrast, we also detected EGFP-positive astroglial cells that were immuno-reactive for glial fibrillary astrocyte protein (GFAP) in the DG and CA1 (**Fig. 4B**). EGFP immunofluorescence was associated with some blood vessels (**Fig. 4C**). This immunofluorescence may be located in endothelial cells, but since it was often closely aligned with that of GFAP, we cannot exclude that EGFP labelling was partly in astrocytic processes surrounding blood vessels (**Fig. 4C**).

**Figure 4:**
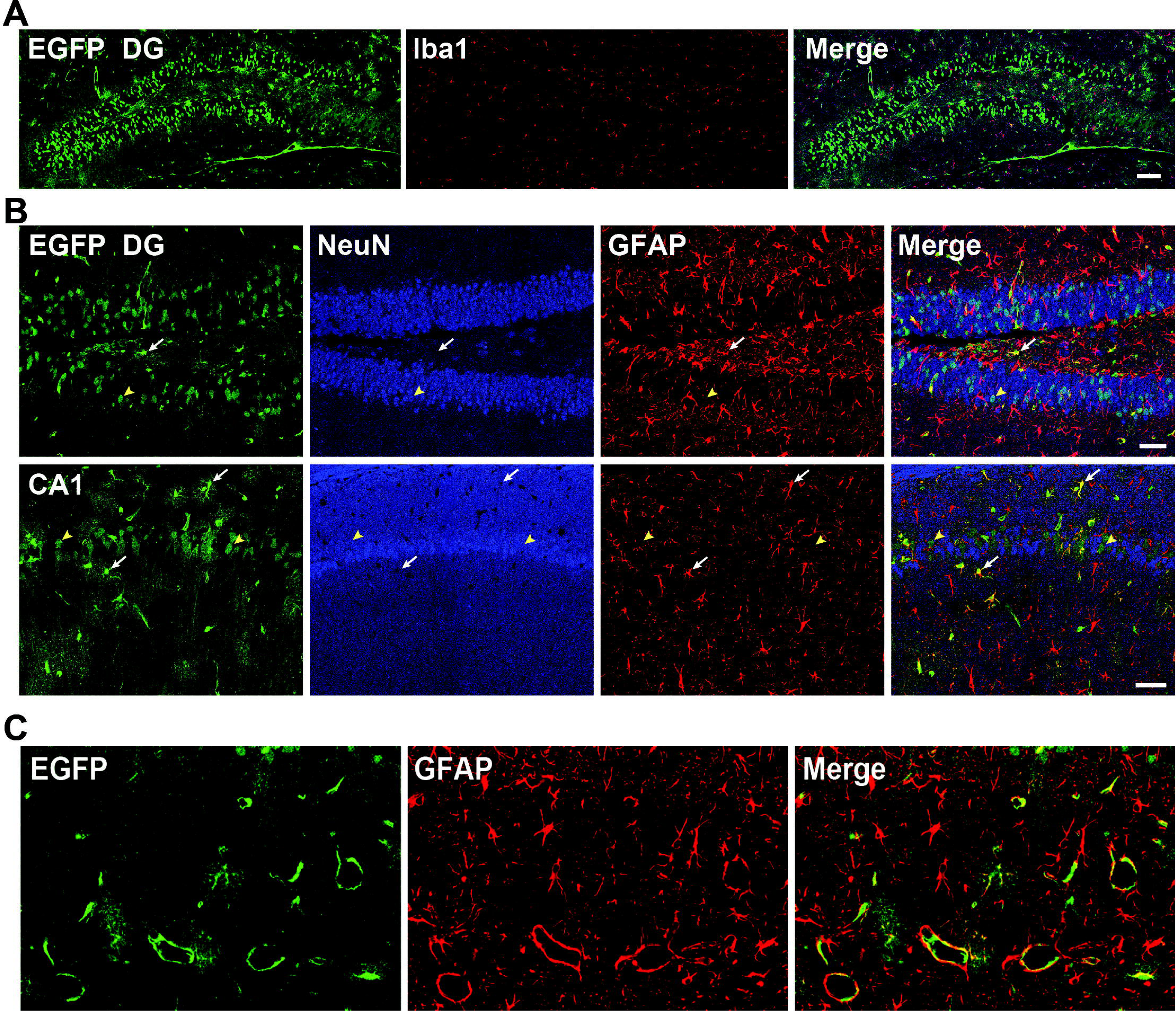
Comparison of glial markers and activity-tagging in hippocampus of Egr1-CreER^T2^ mice after pilocarpine-induced seizures. Egr1-CreER^T2 Tg/+^ x R26^RCE/+^ mice were treated with pilocarpine (70 mg.kg^−1^) and immediately after tamoxifen (120 mg.kg^−1^). One week later, their brain was fixed and brain sections were examined with a confocal microscope for EGFP fluorescence and cell type markers immunofluorescence. **A**) EGFP and IBA1, a marker of microglia, in the DG. Scale bar: 100 µm **B**) EGFP, NeuN, and GFAP in the DG and CA1. Most activity-tagged EGFP-positive cells express the neuron-specific marker, NeuN (yellow arrowhead). Other activity-tagged cells contain the astrocyte-specific marker, GFAP (white arrows). Scale bar: 50 µm **E**) At the junction zone between CA1 and DG, EGFP and GFAP immunolabeling co-localize around blood vessels. Scale bar: 50 µm.

### Appearance and persistence of tagging and comparison with endogenous Egr1 expression

We first determined the time course of EGFP expression in recombinant cells. Egr1-CreERT2 Tg^/+^ x R26RCE^/+^ mice were treated with pilocarpine (70 mg.kg^−1^) and tamoxifen (120 mg.kg^−1^) and they were sacrificed 24, 48, and 72 h later (**Fig. 5A**). We found many cells expressing EGFP as early as 24 h in mice with status epilepticus, and similar labeling in mice sacrificed at 48 h or 72h (**Fig. 5A**). We then assessed the persistence of tagging in the hippocampus of Egr1-CreER^T2 Tg/+^ x R26^RCE/+^ mice one month after pilocarpine (70 mg.kg^−1^) and tamoxifen (100 mg.kg^−1^, ip) treatment. We observed the presence of numerous EGFP-positive cells in all the areas of the hippocampus, showing that the tagging of the neurons was long-lasting (**Fig. 5B, C**). To compare this tagging with spontaneous expression of Egr1 in these mice, we double-stained the sections for Egr1 (**Fig. 5B-E**), knowing that in Egr1-CreER^T2^ mice the BAC promoter drives only the expression of CreER^T2^ and not of Egr1 itself. In most mice, few Egr1-immunopositive cells were found in the DG and CA3 and the EGFP-positive neurons were not Egr1-immunolabeled (**Fig. 5B, C**). In the same mice, many pyramidal neurons were clearly positive for Egr1 in CA1 (**Fig. 5B, C**). However, the EGFP-positive neurons contained variable levels of Egr1 immunoreactivity, including high (**Fig. 5F, arrow**) and absent Egr1 labeling (**Fig. 5F, arrowhead**). In one mouse, we observed a dramatic induction of Egr1 in all the hippocampal areas, including in the DG where the effect was particularly obvious (**Fig. 5E, F**). Since one month after the pilocarpine treatment, the mice are affected by spontaneous recurrent seizures (Curia *et al*., 2008), it is highly probable that this mouse in which Egr1 was highly induced in the hippocampus had seizures just before being killed. In this mouse the EGFP-positive neurons contained high Egr1 immunoreactivity in all the hippocampal areas, including the DG and CA1 (**Fig. 5F**), suggesting that the tagged neurons were also undergoing seizure-induced Egr1-induction. These results show that in Egr1-CreER^T2 Tg/+^ x R26^RCE/+^ mice neuronal tagging is efficient 1 day after the initial induction of seizures in the presence of tamoxifen, that it persists at least a month and that Egr1 can be re-induced in the tagged cells.

**Figure 5:**
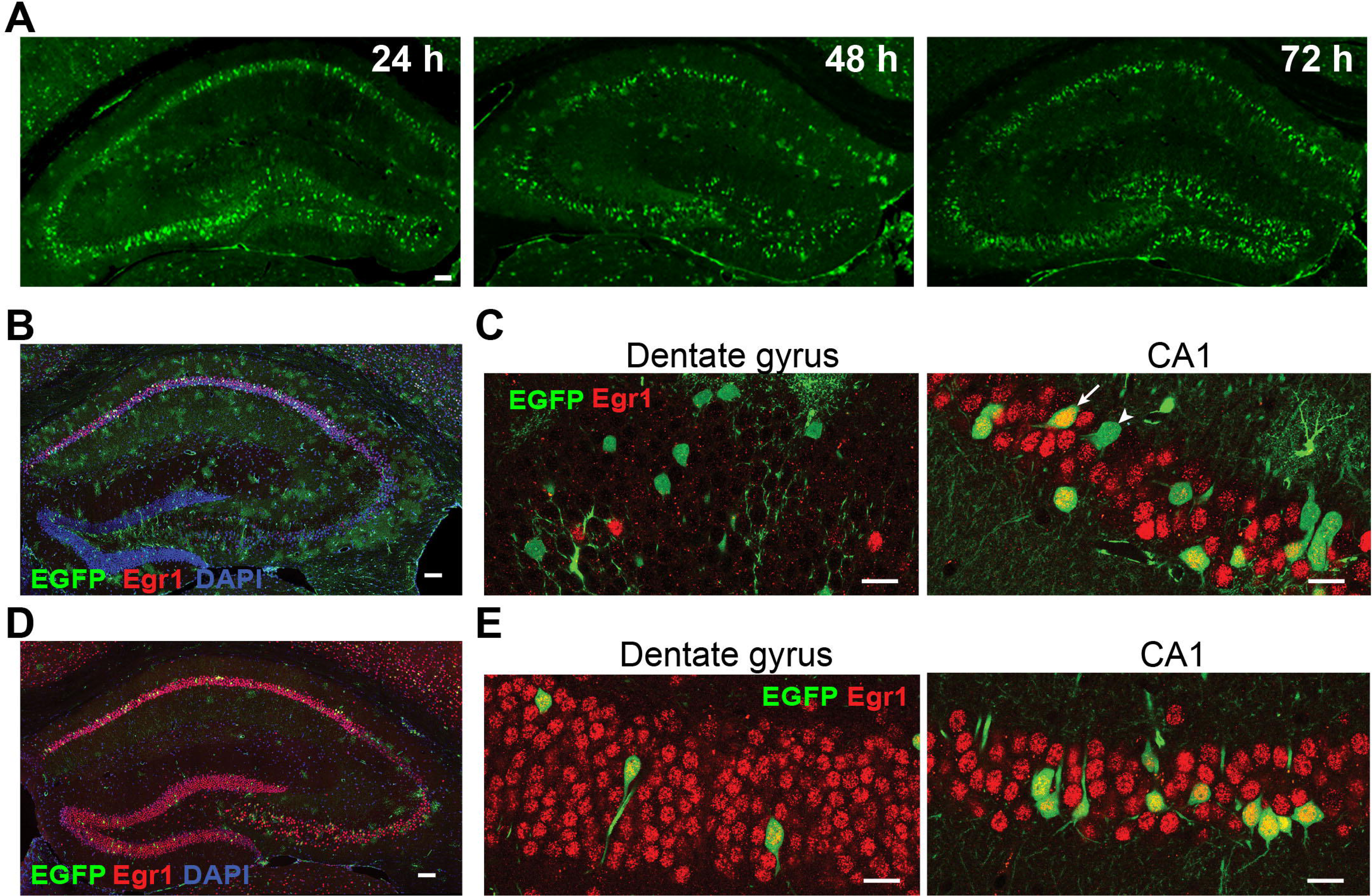
Time-course and persistence of EGFP expression in hippocampus of Egr1-CreER^T2^ mice after pilocarpine-induced seizures and comparison with Egr1. **A)** Egr1-CreER^T2 Tg/+^, R26^RCE/+^ mice were treated with pilocarpine (70 mg.kg^−1^) and tamoxifen (120 mg.kg^−1^) and were perfused for fixation 24, 48 and 72 h later. EGFP-fluorescent cells were observed in all areas of the hippocampus except in a zone between CA1 and CA3 which could correspond to CA2. Many EGFP-positive cells are observed as early as 24 h after the pilocarpine and tamoxifen treatment. Scale bar: 100 µm. **B-C**) Egr1-CreER^T2 Tg/+^ x R26^RCE/+^ mice were treated with pilocarpine and tamoxifen as in **A**. One month later, hippocampal sections were imaged for EGFP fluorescence (green) and Egr1 immunofluorescence (red) with DAPI counter-staining. These images are examples of results observed in most mice, in which the number of Egr1-positive neurons was low in the dentate gyrus. At higher magnification (**C**), EGFP-positive neurons do not express Egr1 in the DG (left panel). In CA1 (right panel), some EGFP-positive neurons express Egr1 (white arrow) and others do not (arrow head). **D-E**) Rare example of a mouse exhibiting intense Egr1 labelling in granule cells of the DG. **E**) At higher magnification, in the DG and CA1 most EGFP-positive neurons express Egr1. Scale bars: **B, D**, 100 µm, **C, E**, 20 µm.

### Activity-dependent genetic labeling of hippocampal cells after pentylenetetrazol-induced seizures

To further explore the possibility to tag activated neurons in Egr1-CreER^T2^ we investigated a different commonly used model of pharmacologically induced epilepsy using pentylenetetrazol (PTZ), a pro-convulsive agent. With this model we tested two reporter lines, R26^Trap^ mice that express EGFP-Rpl10a, and R26^Ai14^ that express tdTomato, both inserted at the Rosa26 locus and Cre-dependent. In addition, since it is not tamoxifen but its metabolite, 4-hydroxytamoxifen (4-OHT), that activates CreER^T2^, we used this compound as a fast-acting activator of Cre (Robinson *et al*., 1991; Guenthner *et al*., 2013). Since 4-OHT can allow a tighter temporal control of target gene recombination than tamoxifen it was important to determine whether it was efficient in our mouse line. In control Egr1-CreER^T2 Tg/+^ x R26^Trap/+^ that were treated with 4-OHT but did not receive PTZ, we observed only a few tagged cells in the CA1 region and the hilus of DG (**Fig. 6A, left panel**). In Egr1-CreER^T2 Tg/+^ x R26^Trap/+^ mice treated with PTZ and 4-OHT, two weeks later a large number of tagged neurons were visible in all regions of the hippocampus, with the exception of CA2 (**Fig. 6A, middle and right panels**). Immunostaining of astrocytes and around blood vessels was low under these conditions. In Egr1-CreER^T2 Tg/+^ x R26^Ai14/+^ mice receiving 4-OHT, we found a fairly high number of tagged cells, even in the absence of PTZ, in CA1 and also in CA3 and DG hilus, but much less in the DG granular layer and in CA2 (**Fig. 6B, left panel**). PTZ treatment dramatically increased the number of tagged neurons in the DG and CA3 (**Fig. 6B, middle and right panels**). Although many tagged pyramidal neurons were observed in the CA1 region, the difference between PTZ-treated and control mice was less obvious than in other hippocampal areas. In all conditions, very few neurons were observed in CA2. These experiments showed that the Egr1-CreER^T2^ system can be used for long-term labeling of neurons that have been activated by epileptic seizures in several models. They also show that these mice respond well to 4-OHT and that the level of baseline recombination, in the absence of specific activation, can be variable depending on the reporter line used.

**Figure 6:**
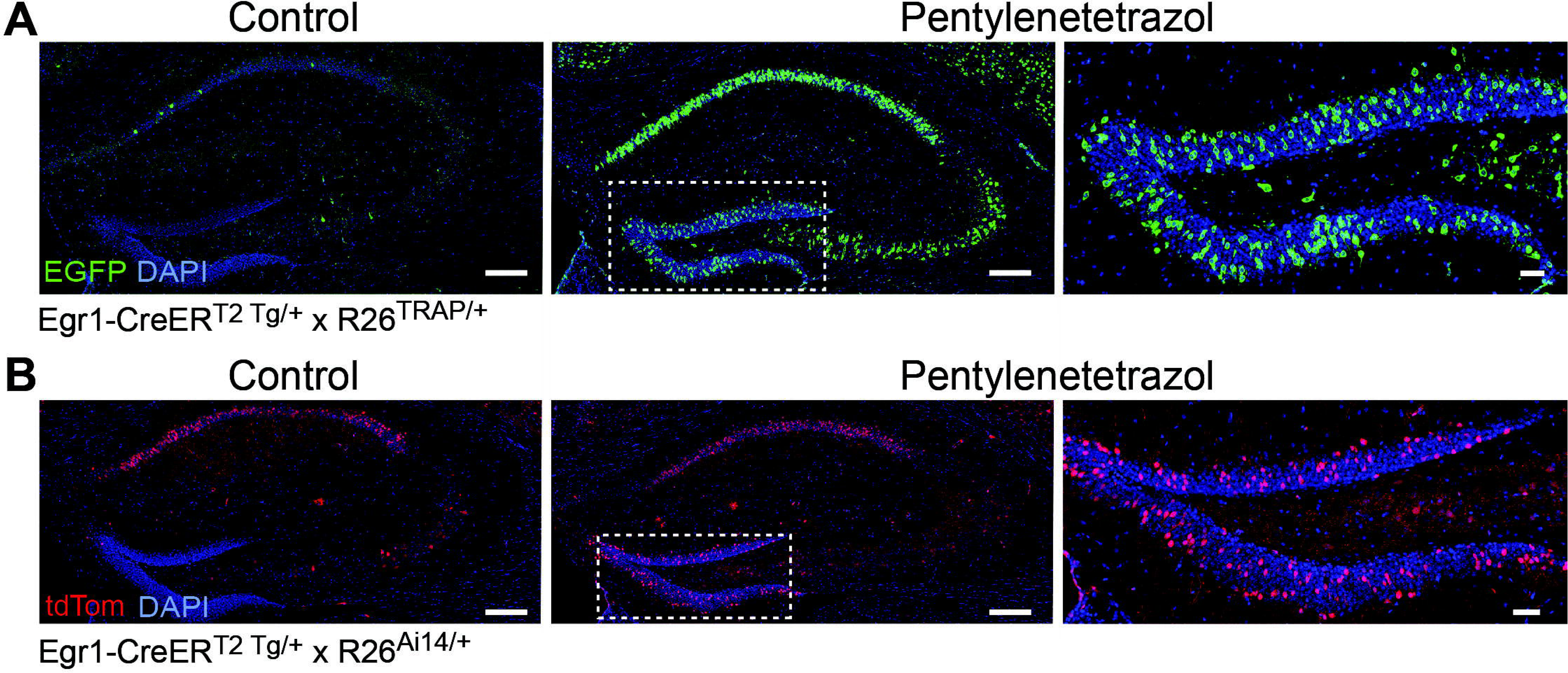
Tagging of hippocampal neurons after pentylenetetrazol-induced epileptic seizures in Egr1-CreER^T2^ mice. Double transgenic mice Egr1-CreER^T2 Tg/+^ and R26^TRAP/+^ (**A**) or R26^RCE/+^ (**B**) were treated with saline (**Control**) or **pentylenetetrazol** (37.5 mg.kg^−1^), and immediately after, with 4-OHT (50 mg.kg^−1^). Two weeks later, sections of their brain were imaged for **EGFP** or tdTomato (**tdTom**) fluorescence and counterstained with **DAPI**. In both mouse lines, numerous EGFP- or tdTomato-tagged neurons are detected in the DG after pentylenetetrazol treatment whereas few tagged neurons are observed in controls. Results in CA1 were variable with either low (**A**) or high (**B**) staining in control conditions. Right panels are higher magnification views of area boxed in middle panel. Scale bar: left and middle, 200 µm; right, 50 µm.

### Activity-dependent genetic labeling of striatal neurons following cocaine administration

Using the Egr1-CreER^T2^ mice, we then switched to a different neuronal model of interest, the activation of striatal neurons by cocaine, a drug that increases extracellular concentration of dopamine and other monoamines and chemically activates brain reward system. Egr1-CreER^T2 Tg/+^ x R26^Ai14/+^ mice were first subjected to a 12-day period of habituation to the experimental environment, manipulations and injections (see **Methods**). Mice were then placed in an open field for 60 min and then i.p. injected with 4-OHT (50 mg.kg^−1^) and immediately after cocaine (20 mg.kg^−1^) or saline (**Fig. 7A**). The animals that received cocaine displayed a marked increase in locomotion, showing that the effects of cocaine were not affected by 4-OHT (**Fig. 7B**). One week after this treatment (**Fig. 7A**), we examined the tagged neurons in the striatum of cocaine- and saline-treated mice (**Fig. 7C**). In saline-treated Egr1-CreER^T2 Tg/+^ x R26^Ai14/+^ mice, we observed a number of tdTomato-tagged cell bodies in the dorsal striatum. They were more numerous in the latero-dorsal part of the dorsal striatum than in the medial and ventral parts. Perivascular labeling was occasionally observed. In cocaine-treated Egr1-CreER^T2 Tg/+^ x R26^Ai14/+^ mice, the number of tagged cell bodies in the dorsal striatum increased markedly (**Fig. 7C**). Quantification indicated that the number of tagged cells was approximately doubled in cocaine-treated mice compared to saline-treated mice (**Fig. 7D**). We compared several reporter mice to determine which combination was optimal to tag striatal neurons following cocaine injection. We noticed that basal tagging, i.e. visible in mice injected with saline, was very different from one line to the other. After an injection of 4-OHT (50 mg.kg^−1^), in saline-treated mice 1.5, 4.8 and 20.8% of the SPNs were tagged in Egr1-CreER^T2 Tg/+^ x R26^RCE/+^, Egr1-CreER^T2 Tg/+^ x R26^Ai14/+^, and Egr1-CreER^T2 Tg/+^ x R26^Trap/+^ mice, respectively (**Fig. 7D**). Cocaine administration significantly increased the number of tagged SPNs in the three reporter lines tested. However, the magnitude of cocaine effects varied depending on the reporter line (**Fig. 7D**). The effect was the strongest in the R26^Ai14^ line (+98%), less in R26^RCE^ mice (+74%) and very low with R26^Trap^ mice (+12%).

**Figure 7:**
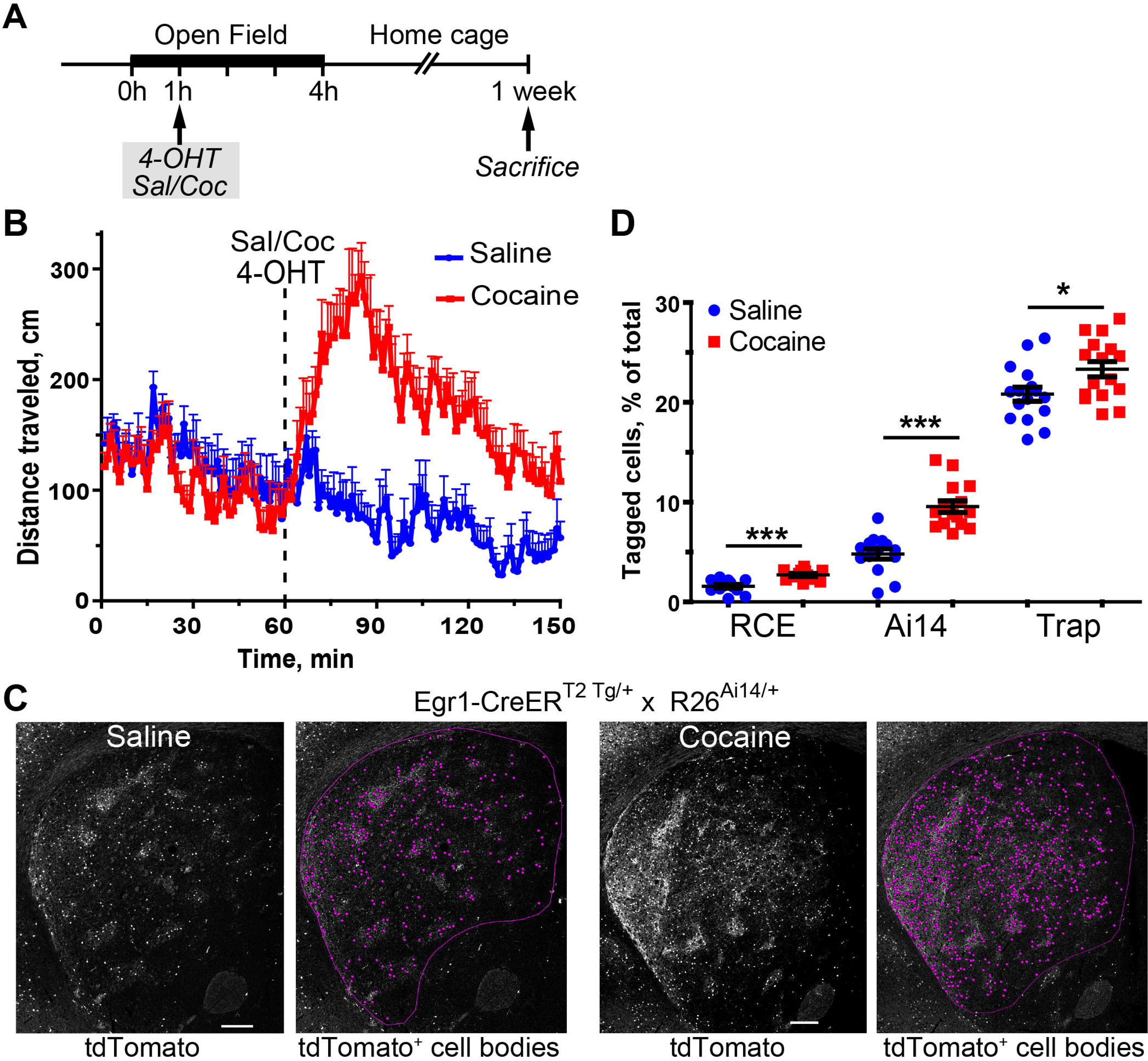
Cocaine treatment increases the number of activity-tagged neurons in the striatum. **A**) Outline of the experimental procedure. **B**) Locomotion in open field of 4-OHT-treated Egr1-CreER^T2 Tg/+^ x R26^Ai14/+^ mice which received immediately after saline or cocaine (20 mg.kg^−1^). Data correspond to means + SEM. Repeated measures two-way ANOVA: effect of treatment, F_(1,13)_ = 34.49, p < 0.01; effect of time, F_(59,767)_ = 2.65, p < 10^−4^; interaction, F_(59,767)_ = 4.08, p < 10^−4^, n = 8 mice per group (see detailed statistical results in Supplementary Table 1). **C**) In Egr1-CreER^T2 Tg/+^ x R26^Ai14/+^ mice, cocaine increased the number of activity-tagged neurons in the striatum, as compared to saline. Sections prepared 1 week after pharmacological treatment as in B, were imaged for tdTomato fluorescence (**left panels**) and positive cell bodies (i.e. tdTomato-tagged cell bodies identified with ImageJ) are indicated by purple dots (**right panels**). Scale bars: 250 µm. **D**) Quantification of the number of activity-tagged neurons in the striatum of R26^RCE^, R26^Ai14^ and R26^Trap^ reporter lines after saline or cocaine treatment. Each point corresponds to one dorsal striatum (2 points per mouse) and means ± SEM are indicated. Results for each mouse line were analyzed by Student’s t test: R26^RCE^, t_(22)_ = 4.59, p = 10^−4^; R26^Ai14^, t_(27)_ = 6.07, p < 10^−4^; R26^Trap^, t_(30)_ = 2.41, p = 0.02 (see detailed statistical results in Supplementary Table 1).

We then characterized the type of cells that were tagged in response to cocaine using an antibody to DARPP-32, a marker of SPNs (Ouimet *et al*., 1984). The tagged cell bodies were virtually all DARPP-32-positive neurons indicating that they were SPNs (**Fig. 8A, B**). We noticed that in both saline and cocaine treated mice tdTomato-positive fibers were unevenly distributed in the striatum, with positive areas that could correspond to striosomes (**Fig. 7C**). To determine the distribution of tagged neurons with respect to the patch-matrix organization of the striatum we used double labeling for calbindin, a marker of matrix SPNs (Gerfen, 1985) (**Fig. 8C**). In contrast with the preferential striosomal localization of tdTomato-positive fibers, a similar frequency of tdTomato tagging of calbindin-positive and - negative cell bodies was observed, which was increased by cocaine (**Fig. 8C, D**). These data indicated that the neuronal tagging induced by cocaine was similar in striosomes and matrix SPNs and suggested that enrichment of tagged fibers in the striosome compartment, observed in this line even in the absence of cocaine, may correspond to cortico-striatal projections towards this compartment. These experiments provide evidence that the Egr1-CreER^T2^ mice provide a good model to specifically label striatal neurons activated by cocaine administration. They also show that the sensitivity and selectivity of the tagging depends on the reporter line.

**Figure 8:**
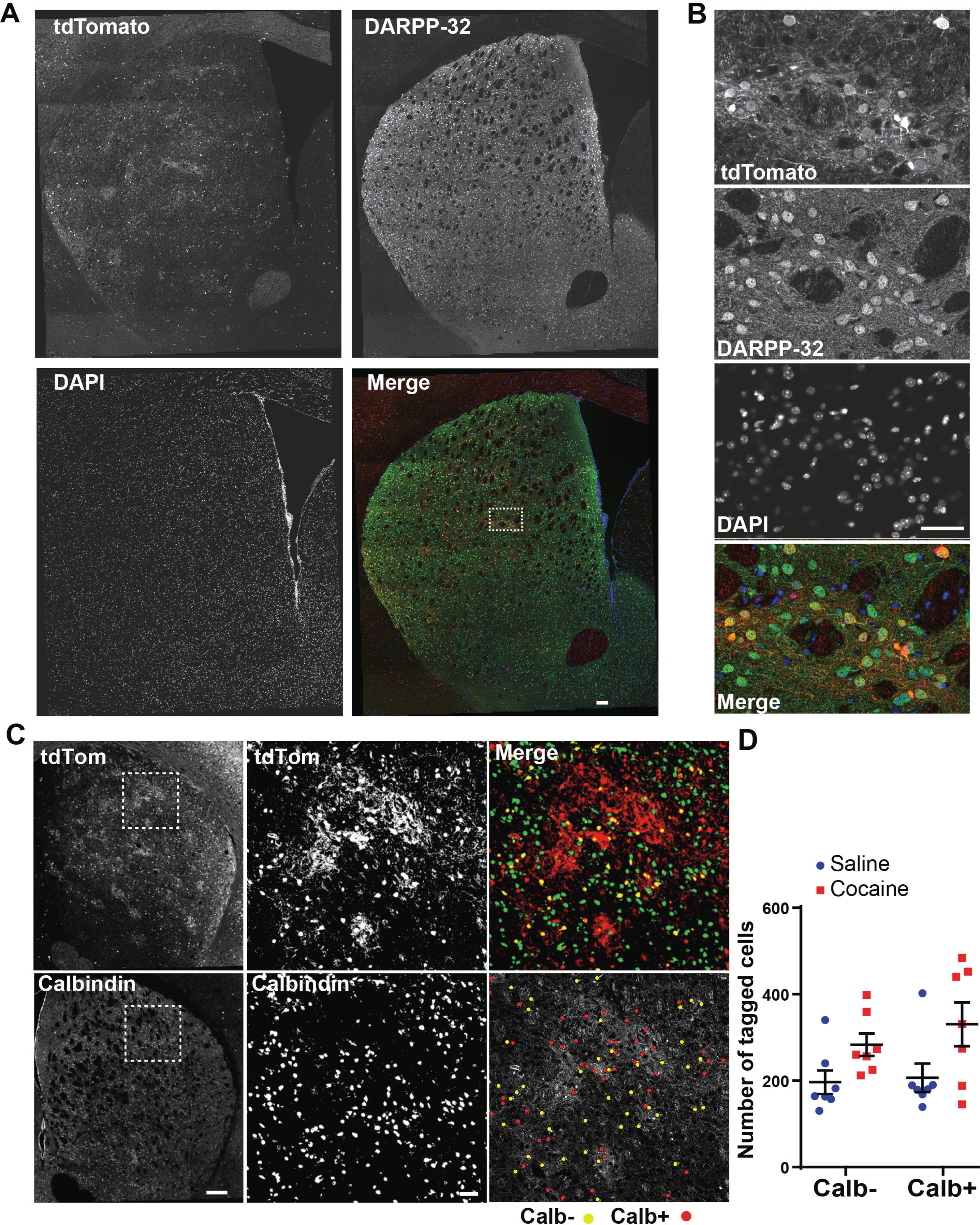
Immunohistological haracterization of activity-tagged neurons in the striatum. **A-B**) Tagged neurons are striatal projection neurons (SPNs). Egr1-CreER^T2 Tg/+^ x R26^Ai14/+^ mice were treated with 4-OHT (50 mg.kg^−1^) and immediately after with cocaine (20 mg.kg^−1^). One week later their brain was processed for confocal imaging of tdTomato fluorescence and DARPP-32 immunofluorescence, a marker of SPNs, and counterstained with DAPI. **A**) Low magnification mosaic pictures of the striatum. **B**) Higher magnification of the area indicated by a dotted rectangle in A. Scale bars, A, 100 µm, B, 50 µm. **C-D**) Cocaine increases activity-tagged neurons in both striosome and matrix compartments in the striatum of Egr1-CreER^T2 Tg/+^ x R26^Ai14/+^ mice. **C**) **Calbindin** immunolabelling and tdTomato (**tdTom**) fluorescence in the striatum of Egr1-CreER^T2 Tg/+^ x R26^Ai14/+^ mouse treated as in A. Middle and right panels are higher magnifications of boxed area in left panels. In the lower right picture, the activity-tagged cell bodies with low and high calbindin immunoreactivities are indicated by red and yellow dots, respectively. The activity-tagged neurons appear widely distributed in both striosomes (low calbindin) and matrix (high calbindin) whereas tdTomato-positive nerve fibers are concentrated in the striosomes. Scale bar: left, 250 µm; middle and right, 50 µm. **D**) Quantification of results as in E. The number of activity-tagged neurons increases after cocaine treatment in both types of neurons with low (Calb-) and high (Calb+) calbindin immunoreactivities. Data are scatter plots with means ± SEM (n = 7 mice per group). Kruskal-Wallis test p<0.05. Mann-Whitney test, effect of cocaine, Calb- p<0.05, Calb+, p= 0.09, pooled Calb-/+, p = 0.003 (see detailed statistical results in Supplementary Table 1).

### Activity-tagged neurons are distributed in the neuronal populations expressing D1 and D2 dopamine receptors

Activity-induced tagging occurred in the striatal projection neurons (SPNs), as indicated by the expression of DARPP-32 within the tagged neurons. In the dorsal striatum, the SPNs are grossly divided into two categories depending on the expression of dopamine D1 or D2 receptors (Drd1 and Drd2, respectively)(Valjent *et al*., 2009). This dichotomy has an important functional relevance since the Drd1- and Drd2-expressing SPNs are the origin of direct and indirect pathways in the basal ganglia network, respectively, which have opposing but coordinated effects (Klaus *et al*., 2019). To determine whether the activity-tagged neurons expressed Drd1 or Drd2, Egr1-CreER^T2 Tg/+^ mice were crossed with Drd1-EGFP ^Tg/+^ (Gong *et al*., 2003) or Drd2-EGFP/Rpl10a^Tg/+^ (Doyle *et al*., 2008) mice that express EGFP fused with Rpl10a, under the control of Drd2 promoter. In the double transgenic mice, we microinjected into the striatum an AAV that Cre-dependently expresses mCherry. Two weeks after the surgery, the mice were treated with cocaine (20 mg.kg^−1^) or saline and immediately after with 4-OHT (50 mg.kg^−1^). Two weeks later, we determined whether the activity-tagged neurons labeled by mCherry also expressed EGFP (**Fig. 9A,B**). A significant increase in the number of mCherry-positive cells in the cocaine-treated mice as compared with saline-treated mice was found in the Drd2-EGFP/Rpl10a^Tg/+^ group, whereas there was only a trend in the other line (**Fig. 9C**). We then determined among the mCherry-tagged cells the proportion of Drd1 or Drd2 expressing cells, by examining the presence of EGFP (**Fig. 9D**). In saline-treated Drd1-EGFP mice, 61% of mCherry-positive neurons were EGFP positive, indicating a slight bias in favor of Drd1-expressing neurons. This result was corroborated by data in Drd2-EGFP/Rpl10a mice in which only 32% of mCherry-tagged neurons expressed EGFP/Rpl10a (**Fig. 9D**). In the cocaine-treated Drd1-EGFP mice, the percent of EGFP-expressing neurons decreased in mCherry-tagged population as compared with saline whereas the reverse trend was observed in the cocaine-treated Drd2-EGFP/Rpl10a mice. The results with cocaine-treated Drd1-EGFP mice indicated a slight bias of mCherry-tagging in neurons expressing Drd2. However, overall the tagged neurons formed a mixed population including neurons expressing Drd1 or Drd2.

**Figure 9:**
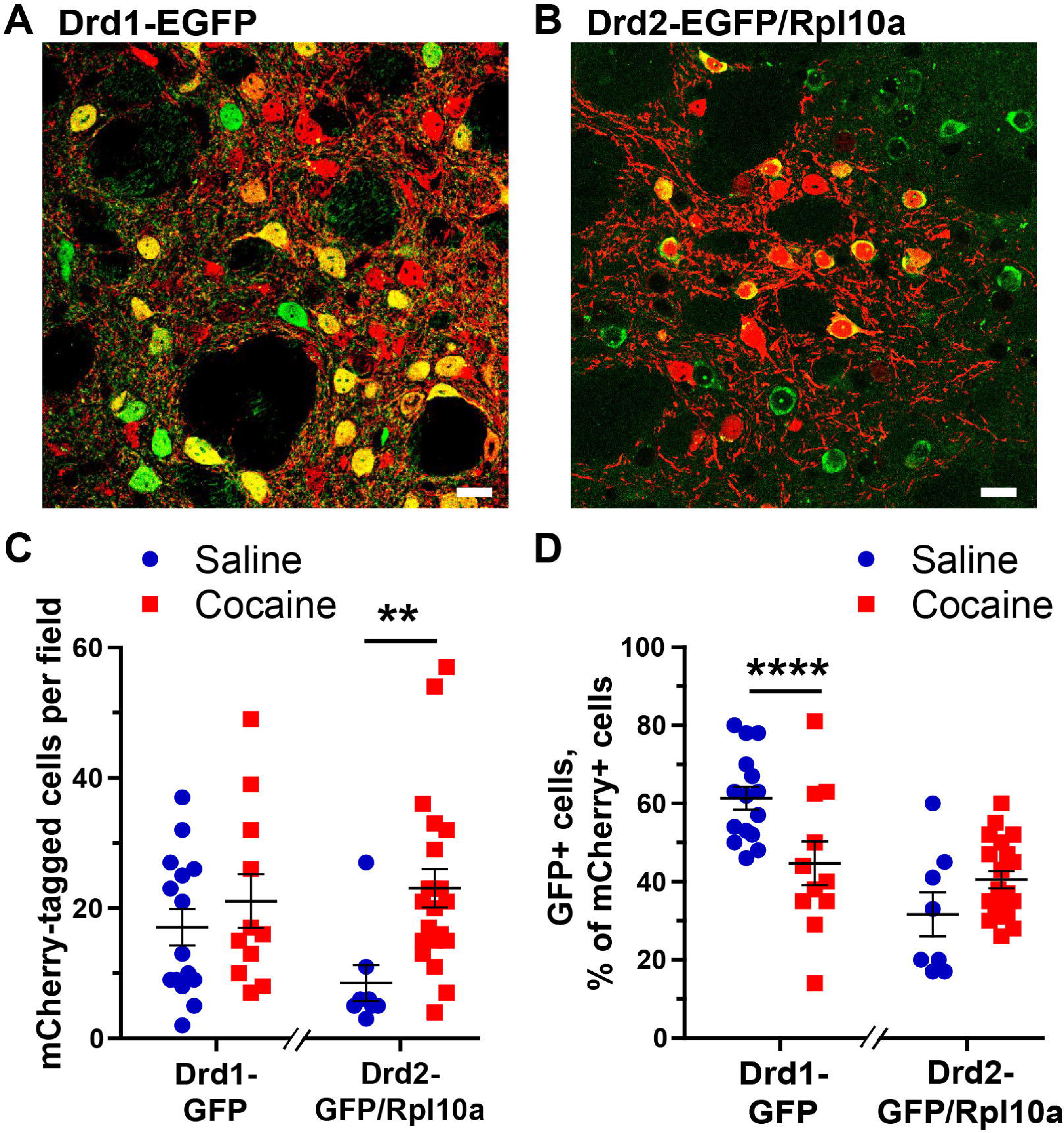
Identification of D1 or D2 dopamine receptors expression in cocaine-induced activity-tagged neurons. Mice carrying the Egr1-CreER^T2^ and R26^Ai14/+^ transgenes and either Drd1-EGFP or Drd2-EGFP-Rpl10a (Drd2^Trap^) transgene were microinjected in the striatum with an AAV that Cre-dependently expresses mCherry. Two weeks later, the mice were treated with 4-OHT (50 mg.kg^−1^) and immediately after with cocaine (20 mg.kg^−1^) or saline. **A**) Confocal image in the dorsal striatum of a Drd1-EGFP mouse 1 week after a cocaine injection (20 mg.kg^−1^) showing mCherry-activity tagged cells (immunodetection, red) and EGFP fluorescence (green) corresponding to Drd1-expressing neurons. **B**) Same as in **A** but in a Drd2^Trap^ mouse. **A** and **B**, scale bar: 20 µm. **C**) Quantification of the number of activity-tagged neurons labeled with mCherry in mice with Drd1-EGFP or Drd2-EGFP/Rpl10a transgene, one week after saline or cocaine injection. Scatter plots with mean ± SEM. Drd1-EGFP, saline, n = 15, cocaine, n = 11, Mann-Whitney test, U = 67.5, p = 0.45; Drd2-EGFP/Rpl10a, saline, n = 8, cocaine, n = 21, Mann-Whitney test, U = 22.5, p = 0.0016 (**, see detailed statistical results in Supplementary Table 1). **D**) Quantification of the percentage of EGFP-positive cells among the mCherry-tagged cells in mice with Drd1-EGFP or Drd2-EGFP-Rpl10a transgene, one week after saline or cocaine injection. Scatter plots with mean ± SEM. Drd1-EGFP, saline, n = 15, cocaine, n = 11; Student’s t test, t_(24)_ = 5.22, p < 10^−4^, ****, Drd2-EGFP-Rpl10a, saline, n = 8, cocaine, n = 20, Student’s t test, t_(26)_ = 0.82, p = 0.42 (see detailed statistical results in Supplementary Table 1).

### Cocaine-induced genetic labeling of striatal neurons depends on ERK activation

The induction of *Egr1* gene by cocaine treatment in striatal neurons is known to be dependent on ERK activation (Valjent *et al*., 2006). We therefore tested whether the tagging induced by cocaine in Egr1-CreER^T2^ mice depended on ERK activation and whether when re-exposed to cocaine a week later ERK was again activated in the neurons tagged after the first injection. With this aim in view, one hour before a first administration of saline or cocaine, Egr1-CreER^T2 Tg/+^ x R26^RCE/+^ mice were pretreated with vehicle or SL327 (50 mg.kg^−1^, i.p.), a brain-penetrant inhibitor of MAPK/ERK kinase (MEK) that prevents ERK activation (see experimental outline in **Fig. 10A**). All these mice also received a 4-OHT injection just before cocaine or saline administration to produce activity-specific recombination events. One week later, the same mice were all sacrificed 15 min after an injection of cocaine to compare the distribution of EGFP tagging and pERK immunoreactivity (**Fig. 10B**). We first compared the number of EGFP-positive neurons in the four groups of mice (**Fig. 10C**). In the vehicle-pretreated mice cocaine injection increased the number of EGFP-tagged neurons in the striatum (**Fig. 10B** and **10C**), as observed above in **Fig. 8D**. In contrast, in mice pretreated with SL327, cocaine did not increase the number of EGFP-tagged neurons (**Fig. 10B** and **10C**). These results show that the cocaine-induced recombination events leading to neuronal tagging depend on ERK activation. Interestingly, we did not detect any effect of SL327 on the number of cells tagged in the absence of cocaine treatment in saline-treated mice animals (**Fig. 10B** and **10C**). This suggests that, in contrast to cocaine-induced tagging, basal tagging of striatal neurons does not result from MEK/ERK activity but from activation of Egr1 expression through different signaling pathways.

**Figure 10:**
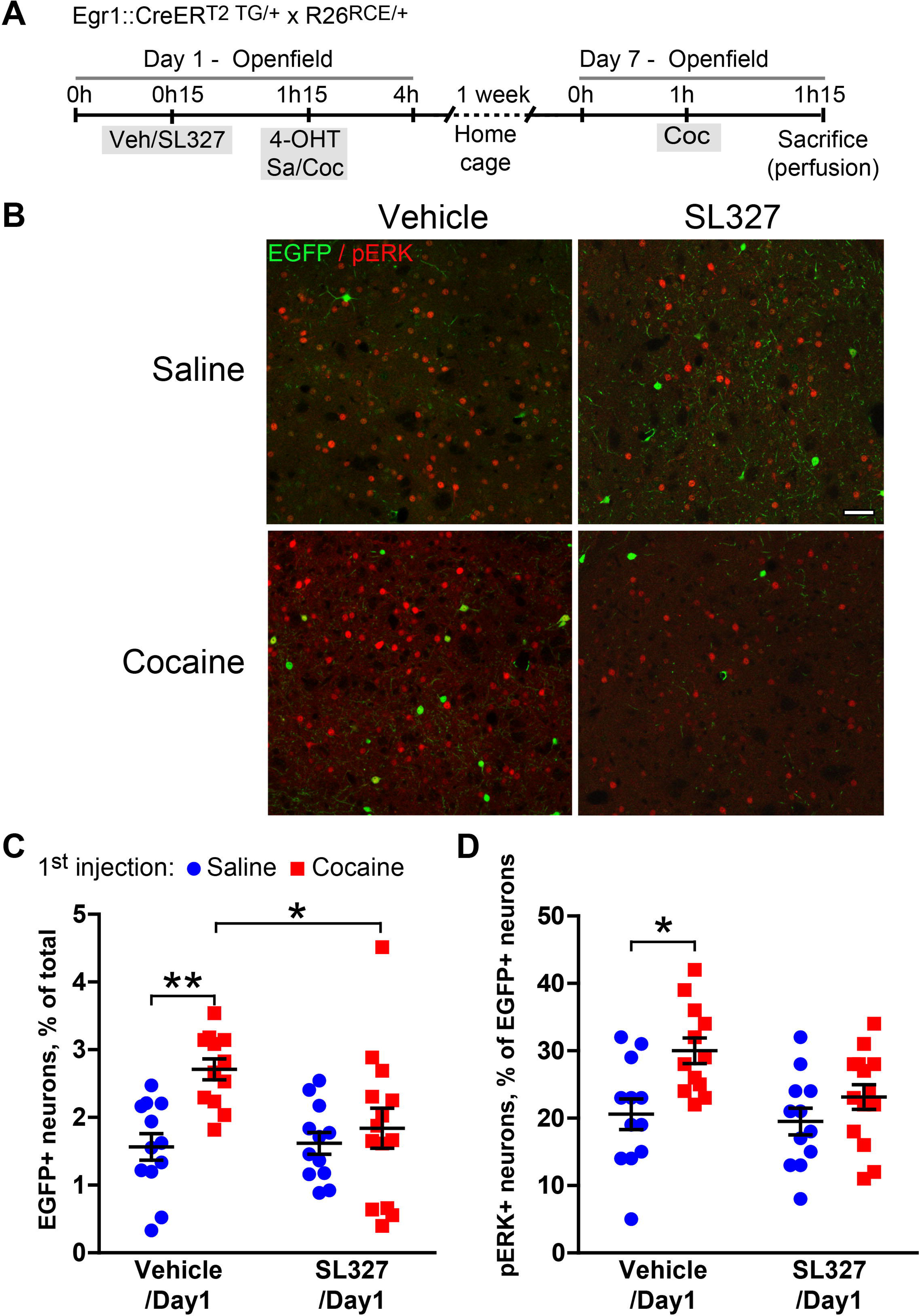
Role of ERK in cocaine-induced neuronal tagging in the striatum. **A**) Outline of the experimental procedure. On Day 1, Egr1-CreER^T2 Tg/+^ x R26^RCE/+^ mice were treated with SL327 (50 mg.kg^−1^) or vehicle, 1 h later with 4-OHT (50 mg.kg^−1^), and immediately after with cocaine (20 mg.kg^−1^) or saline. On Day 7, all the animals received cocaine (20 mg.kg^−1^) and were perfused 15 min later. **B**) Confocal images of the dorsal striatum showing EGFP fluorescence (green) and phospho-ERK (pERK) immunolabeling (red) in the striatum. Scale bar: 50 µm. **C**) Quantification of the number of activity-tagged (EGFP-positive) neurons following the indicated treatments on Day 1. Scatter plots with mean ± SEM, n = 12-14 striata and 6-7 mice per group. Two-way ANOVA: effect of SL327 on Day 1, F_(1, 46)_ = 3.46, NS; effect of cocaine on Day 1, F_(1, 46)_ = 9.66, p = 0.003; interaction, F_(1, 46)_ = 4.40, p = 0.004. Post-hoc Holmes-Sidak’s test, vehicle/saline vs. vehicle/cocaine, p = 0.004, **, vehicle/cocaine vs. SL327/cocaine, p 0.026, *. **D**) Quantification of cocaine-induced pERK immunoreactivity in activity-tagged neurons (i.e. EGFP-labeled). Scatter plots with mean ± SEM. Two-way ANOVA: effect of SL327 on Day 1, F_(1, 46)_ = 3.9, NS; effect of Cocaine on Day 1, F_(1, 46)_ = 10.6, p = 0.002; interaction, F_(1, 46)_ = 2.07, NS. Post-hoc Holmes-Sidak’s test, vehicle/saline vs. vehicle/cocaine, p = 0.013, * (see detailed statistical results in Supplementary Table 1).

These results suggested that the cocaine-tagged neurons were neurons in which ERK was activated by cocaine administration. To test whether ERK was re-activated in the same neurons by a second injection of cocaine, the four groups of mice were treated 1 week later with a cocaine challenge before sacrifice and pERK immunostaining (see experimental outline in **Fig. 10A**). In the population of EGFP-tagged neurons, the number of pERK-positive neurons was higher in the group of animals that had been treated with cocaine during the first test (**Fig. 10D**). Interestingly, if the cocaine-induced ERK activation was blocked by SL327 during the first exposure, this difference was not observed, the percent of pERK-positive neurons remaining similar to that observed in the control animals having received saline injection in the first test. These data show that, in the neurons that are activity-tagged after a first injection of cocaine, ERK is more likely to be activated by a second injection of cocaine. These results therefore suggest that the tagged neurons are part of cocaine-sensitive ensembles.

### Cocaine-tagged neurons are re-activated by re-exposure to cocaine

These results suggested that the neurons tagged by a first injection of cocaine were more prone to be activated again by a re-exposure to cocaine, based on ERK phosphorylation. We then examined whether this was also the case for IEG expression. To test this hypothesis, two groups of Egr1-CreER^T2 Tg/+^ x R26^Ai14/+^ mice were treated on Day 1 with cocaine or saline, right after an administration of 4-OHT (**Fig. 11A**). Seven days later, both groups received cocaine in the same environment (open field) and were sacrificed 90 min later to examine endogenous IEG induction. To control the behavioral effects of cocaine in this experimental setting, locomotor activity was recorded in the open field after the two pharmacological treatments. We observed a sensitization of the locomotor response to the second cocaine injection in the group that already received cocaine the first day (**Fig. 11B**), as previously described (Valjent *et al*., 2010). Both mouse groups were perfused with paraformaldehyde 90 min after the cocaine challenge to analyze Egr1 and Fos expression (**Fig. 11C** and **D**, respectively). As expected, the number of tdTomato-tagged neurons was higher in the mice which had received a first injection of cocaine one week before as compared to those which received saline (**Fig. 11E**). In the two groups treated with cocaine or saline on Day 1, Egr1 immunolabelling was systematically observed in the tdTomato-tagged neurons, suggesting that the tagged neurons had a propensity for Egr1 expression (**Fig. 11C**). However, many neurons with Egr1 immunolabelling were not tagged with tdTomato.

**Figure 11:**
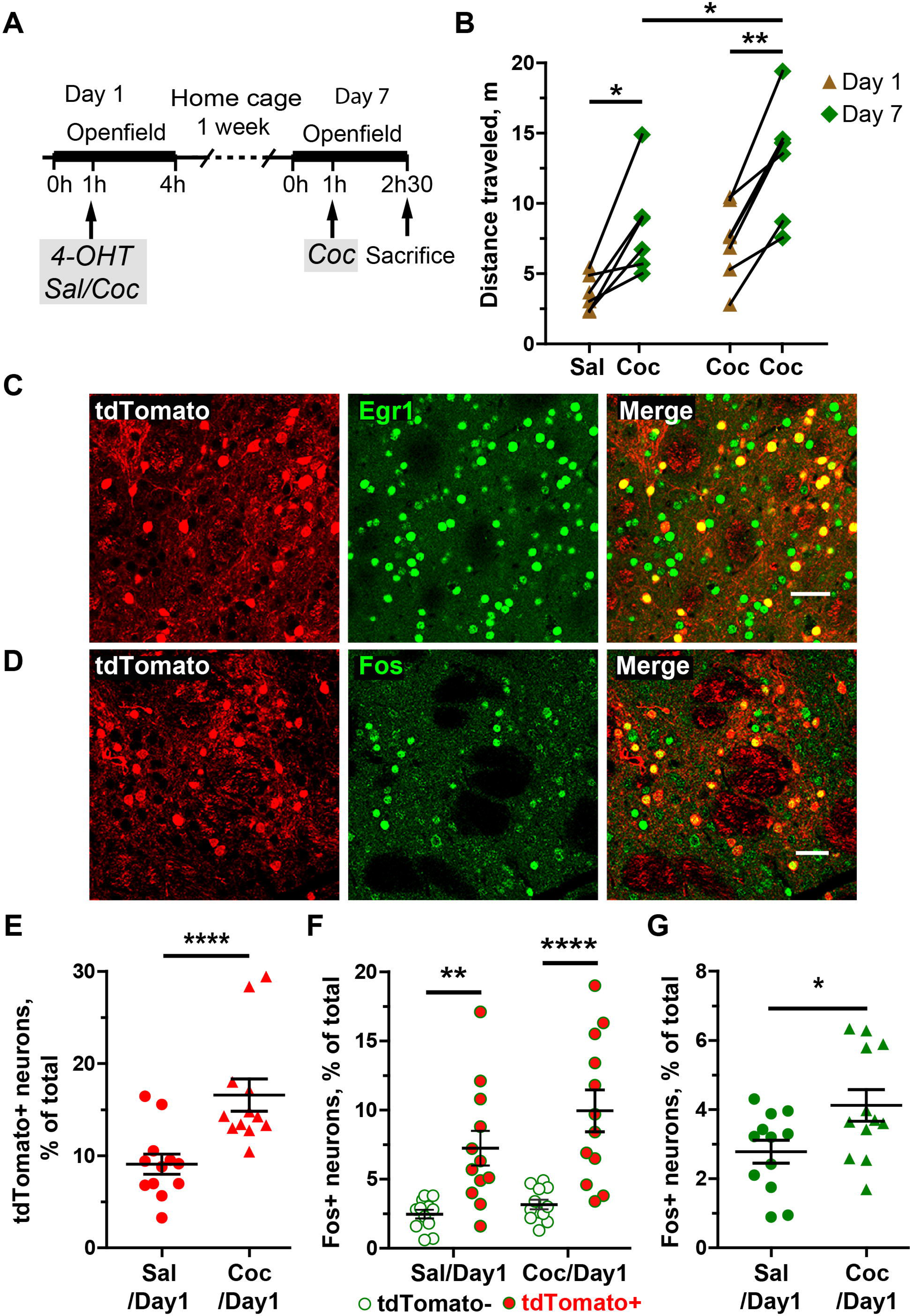
Neurons activity-tagged in response to cocaine are re-activated by re-exposure to cocaine. **A**) Outline of the experimental procedure. On Day 1, Egr1-CreER^T2 Tg/+^ x R26^Ai14/+^ mice were treated with 4-OHT (50 mg.kg^−1^) and immediately after with cocaine (20 mg.kg^−1^) or saline. One week later (Day 7), all the animals received cocaine (20 mg.kg^−1^) and were perfused 75 min later. **B**) Sensitization of the cocaine-induced locomotor activity on Day 7 by cocaine administration on Day 1. Locomotor activity was measured as the distance travelled by the animals during the 45 min after the injection. Paired-scatter plots of responses on Day1 and Day 7. Two-way ANOVA: effect of Day, F_(1,22)_ = 19.2, p = 0.0002; effect of Day 1 treatment (group), F_(1,22)_ = 12.1, p = 0.002; interaction, F_(1,22)_ = 0.22, NS. Post-hoc analysis, Holms-Sidak test, * p < 0.05 (see detailed statistical results in Supplementary Table 1). **C**) tdTomato fluorescence (red) and Egr1 immunolabeling (green) in the dorsal striatum of an animal treated with cocaine on Days 1 and 7. Scale bar: 50 µm **D**) tdTomato fluorescence (red) and Fos immunolabeling (green) in the striatum of an animal treated with cocaine on Days 1 and 7. Scale bar: 50 µm. **E**) Quantification of the number of activity-tagged neurons (tdTomato-positive) in mice treated with saline or cocaine on Day 1, expressed as a % of the total number of neurons. Mann-Whitney test: U = 17, p = 0.0008,. **F**) Quantification of the percentage of Fos-positive neurons in untagged (**tdTomato^−^**) and activity-tagged (**tdTomato^+^)** neurons. Scatter plots with mean ± SEM, two-way ANOVA: effect of tdTomato expression: F_(1,44)_ = 32.7, p < 10^−4^; effect of treatment: F_(1,44)_ = 2.85, NS; interaction: F_(1,44)_ = 0.99, NS. Post-hoc Holmes-Sidak’s test, Tom+ vs Tom-Salc/Coc, p = 0.007, Coc/Coc, p = 10^−4^**. G**) Quantification of the overall number of Fos-positive neurons (Fos+) on Day 7, in mice injected with saline or on Day 1. Student’s t test: t_(22)_ = 2.37, p = 0.027. **B, E-G**, n = 12 striata, 6 mice per group, * p < 0.05, ** p< 0.01, *** p < 0.001. E-G, n = 12 striata (left and right) from 6 mice per group (see detailed statistical results in Supplementary Table 1).

We then investigated whether the tdTomato-tagged neurons were particularly likely to be activated by the cocaine challenge on Day 7 using a different IEG, Fos, as a marker (**Fig. 11D**). The proportion of Fos-positive cells was higher in the tdTomato-tagged neurons than in untagged neurons, in mice treated with cocaine on Day 1 as well as in those which received saline (**Fig. 11F**), indicating that these neurons had a higher responsiveness regardless of the stimulus. When we considered the populations of tagged and untagged neurons separately, the number of Fos+ neurons in each population tended to be higher in the group that received cocaine on Day 1 than in saline-treated mice, but this difference was not significant (**Fig. 11F**). However, the total number of Fos+ neurons (i.e. tagged or untagged) was higher in the mice treated with cocaine on Day 1 than in saline-treated animals (**Fig. 11G**), indicating the existence of a slight hypersensitivity to a second cocaine injection. These results showed that Fos was more easily induced by cocaine challenge on Day 7 in the tagged neurons as compared to untagged neurons, but was similar whether tagging occurred in animals treated on Day 1 with saline or cocaine (**Fig. 11F**). To assess the reactivity of tdTomato neurons we calculated a reactivation coefficient, comparing the number of double-labeled neurons (TdTomato+ and Fos+) to that expected by chance. We divided the proportion of Fos-tdTomato co-labeled neurons among DAPI+ neurons by the chance probability of these neurons to be co-labeled [(Fos+/DAPI+) × (tdTomato/DAPI+)], providing reactivation/chance evaluation (Reijmers *et al*., 2007; Lacagnina *et al*., 2019). In mice treated with cocaine or saline on Day 1 the ratio was > 1, but there was no significant difference between the two groups (saline-treated, ratio = 2.9 ± 0.7, cocaine treated, ratio = 2.4 ± 0.2, mean ± SEM, n = 12 in each group, Mann-Whitney U = 72, p>0.99). This confirmed that tagged neurons were more likely to express Fos after a cocaine test injection on Day 7 than untagged neurons, but that their reactivity was similar whether they had been activated (tagged) in response to cocaine or saline on Day 1.

## Discussion

We have created a new Egr1-CreER^T2^ mouse line allowing to permanently tag neurons in which the *Egr1* gene is induced during specific experimental conditions. The Egr1-CreER^T2^ mice are BAC transgenic mice expressing 4-OHT-activable CreER^T2^ under the *Egr1* promoter dependence. By crossing these mice with several types of Cre-dependent reporter mice, we showed that induction of epileptic seizures in the presence of 4-OHT or tamoxifen induced the appearance of permanently labeled cells in the hippocampus. Similarly, treatment with cocaine in the presence of 4-OHT triggered the labelling of neurons in the dorsal striatum.

### Effective activity-dependent neuronal tagging using Egr1-CreER^T2^ mice

The Egr1-CreER^T2^ mice were created by introducing a recombinant BAC transgene in which the Egr1 coding sequence was replaced by the CreER^T2^ sequence. As a result, the expression of endogenous Egr1 is not modified, as it may be for Fos or Arc in mouse lines used for the activity-tagging of neurons in which the introduction of CreER^T2^ sequence by homologous recombination disrupts the Fos or Arc genes (Guenthner *et al*., 2013; DeNardo *et al*., 2019). This is particularly important in the case of Egr1 since the hemizygous mice for Egr1 gene have a significant behavioral phenotype (Jones *et al*., 2001; Valjent *et al*., 2006; Maroteaux *et al*., 2014).

We crossed Egr1-CreER^T2^ mice with several mouse lines recombined at the Rosa26 locus that Cre-dependently express fluorescent proteins. In general, in the absence of any treatment, the number of fluorescent cells was extremely low in the brain of double-transgenic mice, but it increased significantly when the mice were treated with tamoxifen or 4-OHT. These results clearly showed that in the absence of 4-OHT activation, CreER^T2^ produced by the BAC transgene had no or very low leak activity. However, we noted differences depending on the reporter mouse lines, with a low number of labeled neurons with the R26^Ai14^ line in the absence of tamoxifen or 4-OHT. It is possible that the number of neurons labeled in the absence of treatment increases with the age of the mice, as noted by Guenthner et al. (Guenthner *et al*., 2013). These authors also observed a significant number of tagged cells when the R26^Ai14^ mice were crossed with the Arc^CreERT2^ mice, but not with the Fos^CreERT2^, which express CreER^T2^ under the control of the promoters of *Arc* and *Fos* genes, respectively. This basal labeling could be related to the relatively high basal expression of Arc and Egr1 in some neuronal populations. We also noted that labeled neurons in Arc^CreERT2^ and Egr1-CreER^T2^ mice show distinct regional distributions (Guenthner *et al*., 2013; Denny *et al*., 2014). The differences between the reporter lines is surprising since tagging involves similar recombination events at the same Rosa26 locus in all cases. It was probably unrelated to variations in the level of fluorescent protein expression since for example, the extremely rare recombinant cells in R26^RCE^ mice were intensely fluorescent (see **Fig. 2B**) and the recombination is an all-or-none process. It is more likely that recombination events were less frequent in R26^RCE^ than in R26^Ai14^ mice. We noted that the floxed sequence in the ROSA26 locus is longer in R26^RCE^ than in R26^Ai14^ mice (Madisen *et al*., 2010; Miyoshi *et al*., 2010). It is possible that CreER^T2^ was less efficient at recombining sites in R26^RCE^ than in R26^Ai14^ mice. It is also possible that the genetically modified locus undergoes different epigenetic modifications in the various lines.

We first used an epilepsy model to show that 4-OHT-dependent recombination events in the hippocampus of Egr1-CreER^T2^ mice is directly related to neuronal activity and Egr1 induction in the hippocampus neurons. In all crossing combinations of Egr1-CreER^T2^ mice with reporter mice, the number of tagged cells after tamoxifen or 4-OHT administration was extremely low in most areas of the hippocampus, except in CA1 where Egr1 is expressed in basal condition (Schlingensiepen *et al*., 1991). Although the mice were placed in a new environment and in a normal day-night cycle we did not observe labeled cells in the dentate gyrus, which contrasts with the results reported with Fos^CreERT2^ or Arc^CreERT2^ mice (Guenthner *et al*., 2013; Denny *et al*., 2014; Cazzulino *et al*., 2016). The induction of convulsions by pilocarpine or pentylenetetrazol in the presence of tamoxifen or 4-OHT led to the appearance of activity-tagged cells in the dentate gyrus and CA3 region and a significant increase of their number in the CA1 region. These results are similar to those reported with Fos^CreERT2^ mice following induction of epileptic seizures by pilocarpine or hippocampal electrical stimulations (Dabrowska *et al*., 2019; Kahn *et al*., 2019). These observations are consistent with the very strong induction of *Egr1* in several models of rodent epilepsy (Beckmann & Wilce, 1997). Notably, in our experiments, the number of tagged cells remained low in an intermediate region between CA3 and CA1 presumably corresponding to CA2. Interestingly, CA2 region is best preserved from sclerosis occurring in the Amon’s horn following temporal lobe epilepsy (Steve *et al*., 2014). Our results indicate that this hippocampal area could be less activated than the others following seizure induction. Tagged cells appeared at the earliest time point checked, i.e. 24 h after the induction of epileptic seizure in the presence of 4-OHT, showing that recombination was rapid and led to significant expression of the fluorescent protein in R26^Ai14^ mice. The labeling was stable since many labeled neurons were observed when the animals were examined one month after the induction of epileptic seizures. The number of tagged cells was relatively low, suggesting that recombination events did not occur in all activated neurons. For example, while previous studies indicated Egr1 induction in almost all granular cells of the dentate gyrus after epileptic seizures (Worley *et al*., 1993), at most, only 20% of the granular neurons were tagged when Egr1-CreER^T2^ mice were crossed with R26^RCE^ mice. This indicated that the efficiency of recombination was only partial in the Egr1-expressing neurons and the tagged neurons represent only a fraction of the activated neurons following seizures or other experimental procedures. We also observed tagging of some astrocytes, especially in tamoxifen-treated mice, but less in 4-OHT-treated mice. This labeling may result from astrocytic activation observed in several models of epilepsy, including that with pilocarpine (Seifert *et al*., 2010; Clasadonte *et al*., 2013). The difference between tamoxifen and 4-OHT, which has a shorter duration of action, may indicate a delayed expression of Egr1 promoter in astrocytes.

### Cocaine-induced tagging in the striatum

After 4-OHT administration, we observed in Egr1-CreER^T2^ reporter mice fluorescent tagged neurons a week later in the dorsal striatum and this number increased significantly when the animals were treated with cocaine (20 mg.kg^−1^). Cocaine-tagged neurons were mostly located in the lateral and dorsal regions of the dorsal striatum, and much less in the ventral and medial regions. We noted an absence of tagged cells in the nucleus accumbens. This absence may result from a lower *Egr1* induction by cocaine in this region as observed in previous studies in mice and rats (Moratalla *et al*., 1992; Bhat & Baraban, 1993; Drago *et al*., 1996; Brami-Cherrier *et al*., 2005). These results contrast with those reported with Arc^CreERT2^ mice, which show an increase in activity-tagged neurons in the nucleus accumbens (Ye *et al*., 2016). In non-cocaine-treated mice, we observed a number of tagged neurons regardless of the reporter mice used, probably related to the relatively high level of Egr1 expression in the striatum in baseline condition (Herdegen *et al*., 1995). Their number varied depending on the reporter lines, with the R26^RCE^ mice showing very few tagged cells while more than 20% of striatal neurons were labelled in the R26^TRAP^ line. However, regardless of the baseline level, the number of tagged neurons increased significantly in the striatum of cocaine-treated mice. The best ratio between the numbers of tagged neurons in control and cocaine treatments was obtained with R26^Ai14^ mice, in which cocaine doubled the number of activity-tagged neurons. This increase is similar to that observed with Arc^CreERT2^ mice crossed with R26^Ai14^ (Ye *et al*., 2016). Thus, when crossed with R26^Ai14^ mice, this new Egr1-CreER^T2^ line seems well suited to study neurons after their activation in experimental conditions, especially following cocaine administrations. In contrast, crossing with R26^TRAP^ mice does not appear to be very efficient for EGFP/Rpl10a ribosomal protein expression in cocaine-dependent activity-tagged cells. The number of activity-tagged neurons in this line increases by only 25% in cocaine-treated animals, suggesting that EGFP/Rpl10a -captured mRNAs would arise only to a small extent from cocaine-activated cells and subsequent transcriptomic studies would be inconclusive. These results emphasize that the Egr1-CreER^T2^ system does not allow expression of all effectors in the activity-tagged neurons of interest to the same levels and the reporter mice or AAVs expressing effectors must be thoroughly validated.

Cocaine-induced tagging results from the classical signaling of ERK activation in response to cocaine that depends on the Drd1 and glutamate NMDA receptors as well as DARPP-32 and that results in the *Egr1* gene induction (Valjent *et al*., 2005; Valjent *et al*., 2006). In fact, SL327, a MEK inhibitor preventing ERK activation by cocaine (Valjent *et al*., 2000) completely blocked the cocaine-induced increase in activity-tagged neurons. Interestingly, tagging of neurons in basal condition was unaffected by SL327 treatment suggesting that it results from a basal *Egr1* expression independent of ERK activity.

In the striatum the activity-tagged neurons were exclusively SPNs. By crossing Egr1-CreER^T2^ animals with mice expressing EGFP in either Drd1- or Drd2-SPNs, the activity-tagged neurons were preferentially Drd1-SPNs (about 65%) under basal conditions. In cocaine-treated animals, the number of activity-tagged neurons were more numerous but less frequently Drd1-SPNs and more frequently Drd2-SPNs, suggesting that the cocaine-induced increase preferentially affected D2-SPNs. These results were somewhat surprising, since the induction of *Egr1* in response to cocaine takes place preferentially in Drd1-SPNs (Bertran-Gonzalez *et al*., 2008). However, induction of *Egr1* has been also observed in some Drd2-SPNs and the relationship between *Egr1* induction and neuronal activity-tagging could be complex and possibly biased by the viral infection used in these experiments.

### Sensitivity of tagged cells to reactivation

Using ERK activation and Fos induction as markers, cells tagged by a first injection of cocaine had a higher probability of being activated by a second cocaine injection. It appears that this second activation by cocaine involves more frequently activity-tagged neurons after the first injection of cocaine. We observed that about 10% of activity-tagged neurons were Fos-positive after the second cocaine injection whereas this percentage was only 3 % in the non-tagged neurons. The reactivation clearly exceeded the chance to observe Fos induction in the activity-tagged neurons. However, we observed similar results in controls that received on the first day a saline injection. In these mice, 7% of activity-tagged neurons exhibited Fos induction after the cocaine treatment whereas only 2.5% of non-activated neurons were Fos positive. In this group of mice, the number of Fos-positive or activity-tagged neurons was globally lower, and the total amount of neurons exhibiting both Fos expression and activity-tag was reduced. However, the reactivation/chance ratio was similar to that found in cocaine-pretreated animal. This ratio varied between mice and was not significantly different in saline-pretreated and cocaine-pretreated animals. These observations and the fairly large number of activity-tagged neurons in saline-pretreated mice showed that only a fraction of the activity-tagged neurons is associated with the neural network selectively activated by cocaine. The other neurons could correspond to neurons with significant activity under basal conditions in awake animals or in response to the experimental conditions, such as the introduction of animals into the open-field or the injection. In Arc^CreERT2^ mice, a fairly high activity-tagging is observed in mice in their breeding cage (Guenthner *et al*., 2013; Denny *et al*., 2014; Cazzulino *et al*., 2016) and in Fos^CreERT2^ mice, novel environment increases the activity-tagging of neurons in the nucleus accumbens and striatum (DeNardo & Luo, 2017; DeNardo *et al*., 2019) and other regions (Matos *et al*., 2019). Interestingly, we observed that the neurons that were tagged in the absence of cocaine were also more frequently activated by cocaine injection in the same context. This suggests that the two neuronal populations could correspond to a set of striatal neurons particularly susceptible to respond to cocaine. However, the population of activity-tagged neurons after cocaine treatment was larger than that resulting from context and the number of activity-tagged Fos-positive neurons was higher. Cocaine treatment seems thus to enlarge the striatal neuronal population prone to respond to a second injection of cocaine.

In the DG and CA3 of the hippocampus, activity-tagged neurons during a fear conditioning session are rarely reactivated by the context associated with electric shocks (Denny *et al*., 2014). However, this population of neurons contains a substantial part of the memory of conditioned fear, since its optogenetic inhibition reduces the conditioned response. Similarly, activity-tagged cells in the presence of cocaine could be the basis of responses associated to cocaine. In particular, the Egr1 gene is essential for the place preference conditioned by cocaine and plays an important role in sensitization (Valjent *et al*., 2006). The *Egr1*/activity-tagged neurons could be part of the neural network involved in the long-term effects of cocaine and its identification can lead to a better understanding of their development of lasting alterations linked to cocaine exposure.

In conclusion, the Egr1-CreER^T2^ mouse line provides a novel tool for studying long-lasting tagging of neuronal populations activated in specific conditions during a short time window. The use of 4-OHT to release CreER^T2^ activity provides a better control than tamoxifen. The comparison of various Cre-dependent reporter lines reveals the differences in the number of cells labeled in both basal and stimulated conditions depending on the reporter line. These populations are also partly distinct from those reported in the literature using different immediate-early genes promoters in accordance with the specific regulation of each of them. This underlines that the choice of combination of CreER^T2^ and reporter lines or vectors to activity-label neurons in behaviorally relevant situations has to be adapted to the specific questions addressed. The use of Egr1-CreER^T2^ mouse allowed us to successfully tag relevant populations of neurons in hippocampus and striatum in response to different types of activation. Their characterization opens the way for further investigations.

## Acknowledgements

The present work was supported in part by Inserm and Sorbonne University, by ERC-2009AdG_20090506, FRM DEQ20081213971, and ANR-16-CE16-0018 Epitraces to JAG. Equipment at the IFM was funded in part by *Fondation pour la recherche sur le cerveau* (FRC) and Rotary *Espoir en Tête*, and by DIM *Cerveau et Pensée*, *Région Ile-de-France*. YN was supported by the Fyssen Foundation and KBC by ANR-05-NEUR-0020 and *Région Ile-de-France*. We thank the Mouse Clinical Institute (MCI) for generating the Egr1-CreER^T2^ mice; Drs G Fishell, JC Poncer, P Greengard for providing R26^RCE^, R26^Ai14^, Drd1-EGFP and Drd2-L10aEGFP mouse lines. Relevant experiments were carried out at the IFM Rodent breeding and phenotyping facility and the IFM Cell and Tissue Imaging facility. We are grateful to Dr JC Poncer and N Leroux for advice on protocols of pilocarpine-induced seizures.

## Conflicts of interest

The authors declare no conflict of interest.

## Data Accessibility

Original data as well as non-commercial mouse strain and material are accessible upon request.

## Abbreviations

AAV: adeno-associated virus
BAC: bacterial artificial chromosome
CA1/3: cornu Ammonis area 1/3
CreERT2: Cre recombinase fused to ERT2
DAPI: 4′,6-diamidino-2-phenylindole
DARPP-32: dopamine- and cAMP-regulated phosphoprotein 32,000
DG: dentate gyrus
DMSO: dimethyl sulfoxide
Drd1/2: dopamine D1/2 receptor
ERK: extracellular signal-regulated kinase (a.k.a. MAPK, mitogen-activated protein kinase) pERK, doubly phosphorylated active form of ERK
ERT2: human estrogen receptor with a mutated ligand-binding site restricted to 4-OHT
EGFP: enhanced green fluorescent protein
ES: cells embryonic stem cells
GFAP: glial fibrillary acidic protein
IEG: immediate early gene
KCC2: potassium chloride cotransporter 2
MAPK: mitogen-activated protein kinase (a.k.a. ERK, extracellular signal-regulated kinase)
MEK: MAPK/ERK kinase NS, not significant
4-OHT: 4-hydroxy-tamoxifen
PTZ: pentylenetetrazol
SEM: standard error of the mean
SPN: striatal projection neuron

## Notes

### Competing Interest Statement

The authors have declared no competing interest.

