## Supplemental Table 1 for "Long-lasting tagging of neurons activated by seizures or cocaine administration in Egr1-CreER^T2^ transgenic mice"

Statistical analyses Longueville et al.

### Figure 3B

| Figure | Variable | Group | n mice | n data points | Normality | Mean | SEM | Statistical analysis | Comparison | | Test value | *P* value |
|---|---|---|---|---|---|---|---|---|---|---|---|---|
| 3B | GFP-positive cells (%) | No pilocarpine | 2 | 2 | Small | 0,00 | 0,00 | Kruskal-Wallis test | Overall | | 8,452 | 0,001 |
| | | Pilocarpine no seizure | 3 | 3 | Small | 0,15 | 0,10 | | | | | |
| | | Pilocarpine seizures | 7 | 7 | Small | 6,33 | 2,79 | | | | | |
| | | | | | | | | Dunn's multiple comparisons test | No pilocarpine vs pilocarpine no seizure | | | ns |
| | | | | | | | | | No pilocarpine vs pilocarpine seizures | | | <0,05 |
| | | | | | | | | | Pilocarpine no seizure vs pilocarpine seizures | | | ns |

### Figure 7B, D

| Figure | Variable | Group | n mice | n data points | Normality | Mean | SEM | Statistical analysis | Comparison | DF | Test value | *P* value |
|---|---|---|---|---|---|---|---|---|---|---|---|---|
| 7B | Distance traveled (cm/min) | Saline | 7 | 7 | | | | Two-way ANOVA | Interaction | 59 ; 767 | F = 4,08 | **< 0.0001** |
| | | Cocaine | 8 | 8 | | | | | Time | 59 ; 767 | F = 2,65 | **< 0.0001** |
| | | | | | | | | | Cocaine effect | 1 ; 13 | F = 34,49 | **< 0.0001** |
| 7D | Tagged cells, % of total | RCE Saline | 6 | 12 | Yes | 1,6 | 0,2 | Student's t-test two-tailed unpaired | Saline vs Cocaine | 22 | t = 4.59 | **0.0001** |
| | | RCE Cocaine | 6 | 12 | Yes | 2,7 | 0,2 | | | | | |
| 7D | Tagged cells, % of total | Ai14 Saline | 7 | 14 | Yes | 4,8 | 0,5 | Student's t-test two-tailed unpaired | Saline vs Cocaine | 27 | t = 6,07 | **< 0.0001** |
| | | Ai14 Cocaine | 8 | 15 | Yes | 9,5 | 0,6 | | | | | |
| 7D | Tagged cells, % of total | Trap Saline | 8 | 16 | Yes | 20,8 | 0,7 | Student's t-test two-tailed unpaired | Saline vs Cocaine | 30 | t = 2,41 | **0.02** |
| | | Trap Cocaine | 8 | 16 | Yes | 23,3 | 0,8 | | | | | |

### Figure 8D

| Figure | Variable | Group | n mice | n data points | Normality | Mean | SEM | Statistical analysis | Comparison | DF | Test value | *P* value |
|---|---|---|---|---|---|---|---|---|---|---|---|---|
| 7D | **Number of tagged cells** | Calb- Saline | 7 | 7 | Yes | 196,6 | 27,1 | Kruskal_Wallis test | | 4 groups | 8,57 | <0,05 |
| | | Calb- Cocaine | 7 | 7 | Yes | 283,3 | 26,17 | | | | | |
| | | Calb+ Saline | 7 | 7 | No | 206,7 | 33,2 | | | | | |
| | | Calb + Cocaine | 7 | 7 | Yes | 330,6 | 50,76 | | | | | |
| | | | | | | | | Mann-Whitney | Calb-, Saline vs Cocaine | | 7 | <0,05 |
| | | | | | | | | Mann-Whitney | Calb+, Saline vs Cocaine | | 11 | 0,09 |
| | | Calb-/+ Saline | 14 | 14 | No | 196,6 | 27,1 | Mann-Whitney | Calb-/+, Saline vs Cocaine | 2 groups | 35 | 0,003 |
| | | Calb-/+ Cocaine | 14 | 14 | Yes | 283,3 | 26,17 | | | | | |

### Figure 9C, D

| Figure | Variable | Group | n mice | n data points | Normality | Mean | SEM | Statistical analysis | Comparison | DF | Test value | *P* value |
|---|---|---|---|---|---|---|---|---|---|---|---|---|
| 9C | **Number of mCherry-tagged cell** | Drd1-GFP Saline | 3 | 15 | Yes | 17,1 | 2,8 | Mann-Whitney | Drd1-GFP, Saline vsCocaine | | 67,5 | 0,45 |
| | | Drd1-GFP Cocaine | 3 | 11 | Yes | 21,1 | 4,1 | | | | | |
| | | Drd2-GFP/Rpl10a Saline | 3 | 8 | No | 8,5 | 2,8 | Mann-Whitney | Drd2-GFP/Rpl10a, Saline vs Cocaine | | 22,5 | **0,0016** |
| | | Drd2-GFP/Rpl10a Cocaine | 4 | 21 | No | 23,1 | 3,0 | | | | | |
| 9D | **GFP-positive cells, % of mCherry** | Drd1-GFP Saline | 3 | 15 | Yes | 61,4 | 2,9 | Student's t test | Drd1-GFP, Saline vsCocaine | 21 | 5,22 | **< 0,0001** |
| | | Drd1-GFP Cocaine | 3 | 11 | Yes | 31,6 | 5,6 | | | | | |
| | | Drd2-GFP/Rpl10a Saline | 3 | 8 | Yes | 44,7 | 5,6 | Student's t test | Drd2-GFP/Rpl10a, Saline vs Cocaine | 29 | 0,82 | 0,42 |
| | | Drd2-GFP/Rpl10a Cocaine | 4 | 21 | Yes | 40,5 | 2,2 | | | | | |

### Figure 10

| Figure | Variable | Group | n mice | n data points | Normality test | Mean | SEM | Statistical analysis | Comparison | DF | Test value | *P* value |
|---|---|---|---|---|---|---|---|---|---|---|---|---|
| 10C | EGFP-positive neurons, % of total | Vehicle/Saline | 6 | 12 | Yes | 1,57 | 0,20 | Two-way ANOVA | Interaction | 1 ; 46 | F = 4,40 | **0.004** |
| | | Vehicle/Cocaine | 6 | 12 | Yes | 2,71 | 0,15 | | SL327 effect | 1 ; 46 | F = 3,46 | 0,07 |
| | | SL327/Saline | 6 | 12 | Yes | 1,62 | 0,16 | | Cocaine effect | 1 ; 46 | F = 9,66 | **0,003** |
| | | SL327/Cocaine | 7 | 14 | Yes | 1,84 | 0,30 | | | | | |
| | | | | | | | | Holm Sidak's test | Vehicle, cocaine vs saline | 46 | 3,62 | **0,0044** |
| | | | | | | | | Holm Sidak's test | SL327, cocaine vs saline | 46 | 0,73 | 0,75 |
| | | | | | | | | Holm Sidak's test | SL327 Saline vs. Vehicle Saline | 46 | 0,17 | 0,87 |
| | | | | | | | | Holm Sidak's test | SL327 Cocaine vs. Vehicle Cocaine | 46 | 2,85 | **0,026** |
| 10D | pERK-positive neurons, % of EGFP-positive neurons | Vehicle/Saline | 6 | 12 | Yes | 20,58 | 2,29 | Two-way ANOVA | Interaction | 1 ; 46 | F = 2,07 | 0,16 |
| | | Vehicle/Cocaine | 6 | 12 | Yes | 30,00 | 1,90 | | SL327 effect | 1 ; 46 | F = 3,91 | 0,05 |
| | | SL327/Saline | 6 | 12 | Yes | 19,5 | 1,983 | | Cocaine effect | 1 ; 46 | F = 10,59 | **0,002** |
| | | SL327/Cocaine | 7 | 14 | Yes | 23,14 | 1,842 | | | | | |
| | | | | | | | | Holm Sidak's test | Vehicle, cocaine vs saline | 46 | 3,26 | **0,013** |
| | | | | | | | | Holm Sidak's test | SL327, cocaine vs saline | 46 | 1,31 | 0,99 |
| | | | | | | | | Holm Sidak's test | SL327 Saline vs. Vehicle Saline | 46 | 0,38 | 0,99 |
| | | | | | | | | Holm Sidak's test | SL327 Cocaine vs. Vehicle Cocaine | 46 | 2,46 | 0,11 |

Statistical analyses Longueville et al.

### Figure 11

| Figure | Variable | Group | n mice | n data points | Normality test | Mean | SEM | Statistical analysis | Comparison | DF | Test value | *P* value |
|---|---|---|---|---|---|---|---|---|---|---|---|---|
| 11B | Distance traveled, m | Sal/Coc Day 1 | 6 | 6 | | 3,62 | 0,53 | Two-way ANOVA | Interaction | 1 ; 22 | F = 0,22 | 0,64 |
| | | Sal/Coc Day 7 | 6 | 6 | | 8,38 | 1,47 | | Group (Sal/Coc & Coc/Coc) | 1 ; 22 | F = 12,11 | **0,002** |
| | | Coc/Coc Day 1 | 7 | 7 | | 7,29 | 1,02 | | Injection effect (Day 1 & 7) | 1 ; 22 | F = 19,16 | **0,0002** |
| | | Coc/Coc Day 7 | 7 | 7 | | 13,20 | 1,50 | | | | | |
| | | | | | | | | Holm Sidak's test | Sal/Coc group Day 1 vs 7 | 22 | 2,66 | **0,04** |
| | | | | | | | | Holm Sidak's test | Coc/Coc group Day 1 vs 7 | 22 | 3,57 | **0,009** |
| | | | | | | | | Holm Sidak's test | Day 7 Sal/Coc vs Coc/Coc | 22 | 2,79 | **0,04** |
| 11E | **tdTomato-positive neurons, % of total** | Drd1-GFP Saline | 6 | 12 | Yes | 9,1 | 1,1 | Mann-Whitney | Drd1-GFP, Saline vsCocaine | | 17 | **0,0008** |
| | | Drd1-GFP Cocaine | 6 | 12 | No | 16,6 | 1,8 | | | | | |
| 11F | Fos-positive neurons, % of total | Sal/Coc Tomato- | 6 | 12 | Yes | 2,47 | 0,32 | Two-way ANOVA | Interaction | 1 ; 44 | F = 0,99 | 0,33 |
| | | Sal/Coc Tomato+ | 6 | 12 | Yes | 3,17 | 0,34 | | First inj Sal or Coc | 1 ; 44 | F = 2,85 | 0,098 |
| | | Coc/Coc Tomato- | 6 | 12 | Yes | 7,23 | 1,26 | | Tomato expression | 1 ; 44 | F = 32,71 | **< 0,0001** |
| | | CocCoc Tomato+ | 6 | 12 | Yes | 9,94 | 1,51 | | | | | |
| | | | | | | | | Holm Sidak's test | Salc/Coc Tom+ vs Tom- | 44 | 3,34 | **0,0068** |
| | | | | | | | | Holm Sidak's test | Coc/Coc Tom+ vs Tom- | 44 | 4,75 | **0,0001** |
| | | | | | | | | Holm Sidak's test | Tom- Sal/Coc vs Coc/Coc | 44 | 0,49 | 0,63 |
| | | | | | | | | Holm Sidak's test | Tom+ Sal/Coc vs Coc/Coc | 44 | 1,90 | 0,12 |
| 11G | Fos-positive neurons % of total | Sal/Coc | 6 | 12 | Yes | 2,8 | 0,3 | Student's t-test two-tailed unpaired | Saline vs Cocaine | 22 | t = 2,37 | **0.027** |
| | | Coc/Coc | 6 | 12 | Yes | 4,1 | 0,5 | | | | | |
